# CTCF directly binds G-quadruplex structures to regulate genome topology and gene expression

**DOI:** 10.1101/2025.02.03.636329

**Authors:** Daniela Samaniego-Castruita, Isabella Han, Roxroy C Morgan, Samantha Carpenter, Bryce Williams, Abhijit Chakraborty, Ishwar Radhakrishnan, Ferhat Ay, Samuel A Myers, Anjana Rao, Vipul Shukla

**Affiliations:** Department of Cell and Developmental Biology, Northwestern University, Chicago, USA 60611; Center for Autoimmunity and Inflammation, La Jolla Institute for Immunology, La Jolla, CA 92037; Department of Molecular Biosciences, Northwestern University, Evanston, 60201; Bioinformatics and Systems Biology PhD Program, University of California, San Diego, CA 92093; Department of Pediatrics, University of California San Diego, La Jolla, CA 92093; Laboratory for Immunochemical Circuits, La Jolla Institute for Immunology, La Jolla, USA, 92037; Division of Signaling and Gene Expression, La Jolla Institute for Immunology, La Jolla, USA, 92037; Robert H. Lurie Comprehensive Cancer Center, Northwestern University, Chicago, USA 60611; Center for Human Immunobiology, Northwestern University, Chicago, USA 60611

**Author notes:** These authors contributed equally (Daniela Samaniego-Castruita, Isabella Han, Roxroy C Morgan). Co-Lead authors (Anjana Rao, Vipul Shukla).

## Abstract

DNA G-quadruplexes (G4s) are non-B form secondary DNA structures that are highly conserved across evolution. G4 structures occupy key regulatory sites in the mammalian genomes and are implicated in several cellular processes. However, the mechanisms by which G4s contribute to distinct facets of genome function are not well understood. Here, we conduct a proteomics screen with G4s of diverse topologies to uncover novel G4 binding activities in genomic regulators of nucleosome remodeling, paraspeckle assembly, RNA splicing, and 3D genome organization. Among the most prominent hits, we identify the genomic architectural protein, CTCF, as one of the strongest G4 binders. Building on this discovery, we perform extensive biochemical validation of CTCF-G4 interaction and identify CTCF mutants, with pronounced affinity for G4s over its consensus DNA motif. By implementing well-established approaches and developing new G4 mapping tools, we define a comprehensive catalog of genomic G4s and investigate their association with CTCF binding. Using genetic reconstitution of mouse embryonic stem cells with a G4-specific CTCF mutant, we ascribe genomic functions to CTCF-G4 interactions in regulation of CTCF occupancy, chromatin looping and gene expression. Interestingly, our studies reveal a subset of G4-linked chromatin loop anchors that form “persistent loops” which are retained even upon CTCF depletion. Collectively, our work establishes the G4 binding activity of CTCF and provides new insights into the functional significance of G4 structures.

## Introduction

G-quadruplexes (G4s) are non-B-form DNA secondary structures that form at guanine-rich regions in the genome^1–7^. G4s form when four stretches of guanine (G) bases within G4 sequence motifs (G_≥3_N_1-7_G_≥3_N_1-7_G_≥3_N_1-7_G_≥3_) come together in planar symmetry, stabilized by atypical Hoogsteen hydrogen bonding and monovalent cations (typically K^+^ ions)^2–5,8^. Depending on the sequence context, relative orientation of the DNA strands, and the spacing within guanine stretches, G4s can adopt distinct topologies^5^. Notably, G4s are highly conserved across evolution^7,9^, and putative G4-forming sequences (pG4s) are enriched at gene promoters^4,10^, 5’ untranslated regions^4,11^, DNA replication origins^4,12^ and telomeres^4–6^, indicating that these structures likely have important functions in the genome. Consistent with this notion, studies have uncovered the association of G4 structures with regulation of gene expression^2,4,5^, chromatin accessibility^13^, DNA methylation^14,15^, genomic stability^15–17^, telomere homeostasis^4,5,18^ and three-dimensional (3D) genomic architecture^19–21^. That said, the mechanisms by which G4s regulate these diverse genomic processes continues to be an area of active investigation. Recent affinity-based and proximity-based labelling approaches have identified G4 binding activity in transcription factors (SP1, FUS, E2F and NRF1)^22^ as well as protein complexes involved in regulation of DNA methylation (DNMT1, UHRF1) and chromatin accessibility (<u>SWI</u>tch <u>s</u>ucrose <u>n</u>on-<u>f</u>ermentable, SWI/SNF complex)^23^. However, the biochemical basis of G4 interaction with these proteins, as well as the genomic contexts in which they occur, warrants additional investigation to functionally elucidate G4 biology.

Among the genomic functions of G4s, recent reports have proposed an intriguing link between G4 structures and higher-order genomic folding^19–21,24^. Mammalian genomes are segregated into chromosomal territories that are further partitioned into distinct compartments, separating gene-rich active regions (compartment A) from the gene-poor inactive regions (compartment B)^25–30^. At the scale of tens to hundreds of kilobases, the genome is further assembled into self-interacting regions known as topologically associating domains (TADs), with higher propensity of genomic interactions observed within, rather than, between TADs^25–29^. The TAD structures mediate enhancer-promoter interactions and are important for regulating cell-type specific gene expression programs^25–29^. Interestingly, G4 structures are shown to be enriched at TAD boundaries and are implicated in regulating insulation strength of TADs^19^. On a molecular level, mammalian TAD boundaries are frequently associated with binding of CCCTC-binding factor (CTCF) and the cohesin complex^28,30^. CTCF is a highly conserved Zinc-finger (ZF) containing DNA binding protein that is proposed to play a central role in a linear tracking model known as “loop-extrusion” for long-range chromatin interactions^26,31–34^. The loop-extrusion model posits that CTCF, bound to its DNA consensus motifs, particularly in convergent orientation, impedes the movement of the cohesin ring complex to define anchors of chromatin loops and TAD boundaries^30,33,35^. Though which convergent CTCF sites demarcate loop anchors or TAD boundaries and if additional determinants are required for specifying the boundary elements for chromatin looping are open questions.

The binding of CTCF to chromatin is highly dynamic with residence times in the range of minutes, although once associated to certain sites, CTCF can remain bound for tens of minutes^36,37^. CTCF binding in the genome is most well-characterized at a 12-15 base pair DNA consensus motifs (5--NCANNA-G(G/A)N-GGC-(G/A)(C/G)(T/C)-3’) which are present at ∼80% of all CTCF binding sites^32,34,38,39^. CTCF harbors 11 cysteine2-histidine2 (C2H2)-type ZFs and structural studies have shown that the central ZFs 4-7 are critical for recognizing the consensus DNA motif^26,32,38,39^. It is estimated that in any given cell type roughly 30-60% of CTCF consensus motifs are occupied by CTCF^40^. It is worth mentioning that the affinity of CTCF for different genomic DNA motifs can differ greatly and the mechanisms for selective binding of CTCF to specific sites are not fully understood^39,41^. In this regard, a recent study suggested that G4 structures can enhance binding of CTCF to its consensus DNA sequence particularly at weaker motifs^20^. However, the authors in this study could not detect direct interaction of CTCF with G4s^20^. Moreover, the structural basis and the biophysical principle for the proposed model, which would require precise positioning of G4s relative to the CTCF consensus motifs, are unclear^20^. The relationship between CTCF and G4s is further complicated by another recent report, which proposed both direct and indirect roles of G4s in CTCF recruitment^24^. Besides its binding to DNA, CTCF is also reported to bind chromatin-associated RNA^42,43^. Taken together, these recent studies highlight that CTCF occupancy in the genome is more nuanced than previously appreciated.

Here, by performing a proteomics screen with G4s of distinct topologies, we uncover G4 binding activity in several known and novel proteins that play key roles in genome biology. We identify G4 binding activity in protein complexes involved in nucleosome remodeling, paraspeckle assembly, RNA splicing, and 3D genome organization. Among the major hits from the screen, we report robust G4 recognition by CTCF. We perform biochemical validation of CTCF-G4 interaction and characterize specific CTCF mutants with strong binding preference for G4s. Furthermore, we utilize murine embryonic stem cells (mESCs) with degron-tagged CTCF to acutely deplete endogenous CTCF and reconstitute with a G4-specific CTCF mutant. Using these experimental designs, we dissect the genomic roles of CTCF-G4 interaction for regulation of chromatin looping and gene expression. Together our work provides compelling evidence for CTCF-G4 interaction and contributes to the emerging roles of G4 structures in genome function and organization.

## Results

### A proteomics screen identifies CTCF as a G4 binding factor

To investigate the nuclear functions of DNA G4 structures, we used four biologically relevant oligonucleotides (oligos) which form G4s of parallel (Kit1 and Kit2), anti-parallel (Spb1) and mixed propellor/intermolecular (Telomere: Telo) topologies (**Fig.1A**). We minimally mutated the G-stretches in each of these oligos to generate respective non-G4 control oligos (**Fig.1A**). To control for potential differences in biotinylation that may in turn impact pulldown efficiencies, we extended the G4 and non-G4 oligos to carry a complementary DNA sequence, which was annealed to a common biotinylated adaptor oligo (**Fig.1A**). This oligo design also provides the double strand DNA (dsDNA)-G4 junction associated with G4s *in vivo* (**Fig.1A**). We prepared nuclear lysates from primary murine B cells stimulated with lipopolysaccharide (LPS) and interleukin-4 (IL-4) for 72 hours and used a B cell lymphoma cell line CH12F3 (CH12) cells. Since G4 formation is sensitive to salt concentrations, we used hypotonic nuclear lysis combined with dounce homogenization and performed buffer exchange to replace the lysis buffer with an intracellular salt buffer (110 mM KCl and 10.5 mM NaCl), to mimic conditions conducive for G4 formation in cells (**Fig.1A**). For pulldown assays, we performed in-solution interaction of biotinylated G4 and non-G4 oligos with nuclear lysates and captured bound-fractions on streptavidin conjugated magnetic beads. For initial tests, we added G4-specific Flag-tagged single chain variable fragment BG4 (BG4) antibody^44^ in nuclear lysates and observed robust enrichment of Flag-BG4 and a well-known G4 helicase, Bloom (BLM) with the G4 oligos compared to non-G4 oligos (**Fig.1B**).

**Figure 1:**
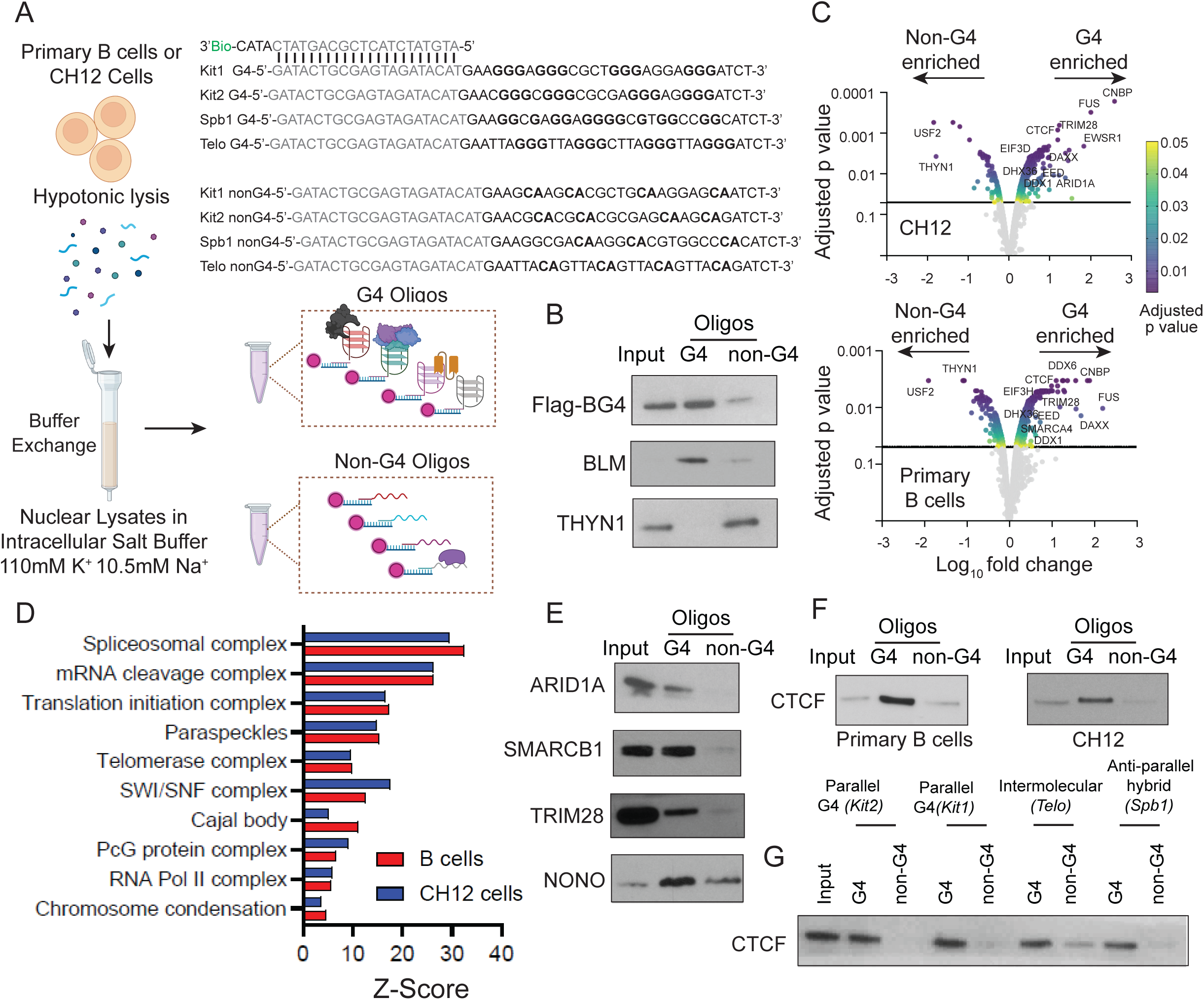
Proteomics screen for G4 binding proteins A. A schematic illustrating hypotonic lysis of primary B cells or CH12 cells combined with dounce homogenization and buffer exchange, followed by incubation with biotinylated G4 or non-G4 oligos. Top right: nucleotide sequence of each oligo used, gray indicating sequence complementary to the biotinylated (Bio) adaptor anchor, and bolded letters indicating guanine stretches (G4 oligo) for G4 formation or mutation of guanine stretches in non-G4 oligos. B. Immunoblot showing the enrichment of Flag-BG4 (spike-in) and a G4 helicase BLM in nuclear lysates pulled down with G4 or non-G4 oligos. THYN1 is included as ssDNA binding protein used as negative control. C. Enrichment profile of G4 protein pulldown fractions from CH12 (top) and primary B (bottom) cells quantified by Tandem Mass Tagging followed by Mass-spectrometry. D. Top protein complexes enriched with G4s from the proteomics screen in primary B cells (red) or CH12 cells (blue). Protein complex enrichment was determined by Metascape analysis with proteins showing 2-fold increase over non-G4 controls and p-adjusted value >0.05. The X-axis represents the Z-score. E. Immunoblot showing the enrichment of ARID1A, SMARCB1, NONO, and TRIM28 in G4 pulldown compared with non-G4 control oligos. F. Immunoblot showing the enrichment of CTCF upon pull down with G4 oligos or non-G4 control oligos from primary B cells (left) or CH12 cells (right). G. Immunoblot showing the enrichment of CTCF with G4 oligos of distinct topologies, including parallel (Kit2), parallel-propeller (Kit1), intermolecular (Telo), and anti-parallel hybrid (Spb1) G4. The relative enrichment is compared to respective control oligos. Data is panels B, E, F and G are representative of at least three independent experiments.

To discover a catalog of G4-interacting proteins, we subjected the G4 and non-G4 pulldown fractions (two biological replicates from primary B and CH12 cells) to quantitative, mass spectrometry-based proteomics (**Fig.1C, Sup.Fig.1A-B**). Compared with non-G4 control oligos, our results showed G4-mediated enrichment of several protein complexes involved in important genomic functions, such as RNA splicing, mRNA cleavage, telomere homeostasis, nucleosome remodeling (SWItch-Sucrose Non-Fermentable or SWI/SNF complex), chromosome condensation (including CCCTC-binding factor or CTCF) and histone modifications (Polycomb repressor or PcG complex) (**Fig.1D, Sup.Fig.1C**). In fact, several subunits of the spliceosomal, mRNA cleavage, SWI/SNF, PcG and paraspeckle assembly were enriched in G4-bound fractions (**Sup.Fig.1C**). Importantly, many of the hits from our screen overlapped proteins that were recently identified using an analogous approach labeling G4 interacting proteins with click chemistry using a G4 ligand Pyridostatin (PDS)^23^; these common hits included proteins involved in nucleosome remodeling, RNA splicing and mRNA processing, among others (**Fig.1D**). Even though we used DNA G4s, we also observed enrichment of RNA processes such as translation initiation, where G4 structures have been previously implicated^4,45^. We also noted G4 binding activity in several proteins associated with paraspeckles and Cajal bodies, which are membrane-less nuclear bodies with biomolecular condensate-like features. We corroborated some of the top targets using western blotting and confirmed the enrichment of SWI/SNF subunits ARID1A and SMARCB1, heterochromatin associated protein TRIM28 (Tripartite motif-containing 28), the paraspeckle associated protein NONO (Non-POU domain-containing octamer-binding protein) (**Fig.1E**), and CTCF in G4-bound fraction compared with non-G4 control oligos (**Fig.1F**).

Since many of the identified proteins are subunits of multimeric protein complexes, we focused on characterizing the G4 binding activity of CTCF, which directly binds DNA and was recently shown to associate with G4 structures^20,24^. To begin, we tested the G4 binding specificity of CTCF to the four G4 oligos used in our screen. Compared to the respective non-G4 controls, CTCF showed strong binding to each of the four G4s of different topologies, highlighting that CTCF recognizes G4 structures rather than an underlying DNA sequence within the G4 oligos (**Fig.1G**). Altogether, the proteomics screen uncovered the ability of several important regulators of genomic function, including the genomic architectural protein, CTCF, to bind G4 structures specifically.

### Biochemical characterization of CTCF-G4 interaction

For defining the biochemical basis of G4 recognition by CTCF, we established a system for *in vitro* transcription and translation of CTCF using T7 polymerase and rabbit reticulocyte lysate (**Fig.2A**). *In vitro* synthesized full length (FL) CTCF showed strong binding to biotinylated G4 oligos compared to the corresponding non-G4 oligos (**Fig.2A**). To further characterize the specific domains of CTCF directly involved in G4 recognition, we performed deletional mutagenesis. CTCF is composed of 11 C2H2-type zinc fingers (ZFs) and unstructured regions at N– and C-termini. We found that the N– and C-terminal regions, as well as ZFs 1-3 and ZFs 8-11 are not essential for G4 recognition by CTCF (**Fig.2A**). However, akin to its binding to the consensus dsDNA motif, the central ZFs 4-7 (117 amino acids) were both essential and sufficient for G4 recognition (**Fig.2A**). To investigate this binding further, we generated the recombinant CTCF with central ZF4-7 fragment using bacterial expression (**Fig.2B**). The CTCF ZF4-7 fragment displayed clear binding to G4 DNA compared to non-G4 controls (**Fig.2C**), ruling out the possibility that CTCF ZF 4-7 interacts indirectly with G4s through a binding partner.

**Figure 2.**
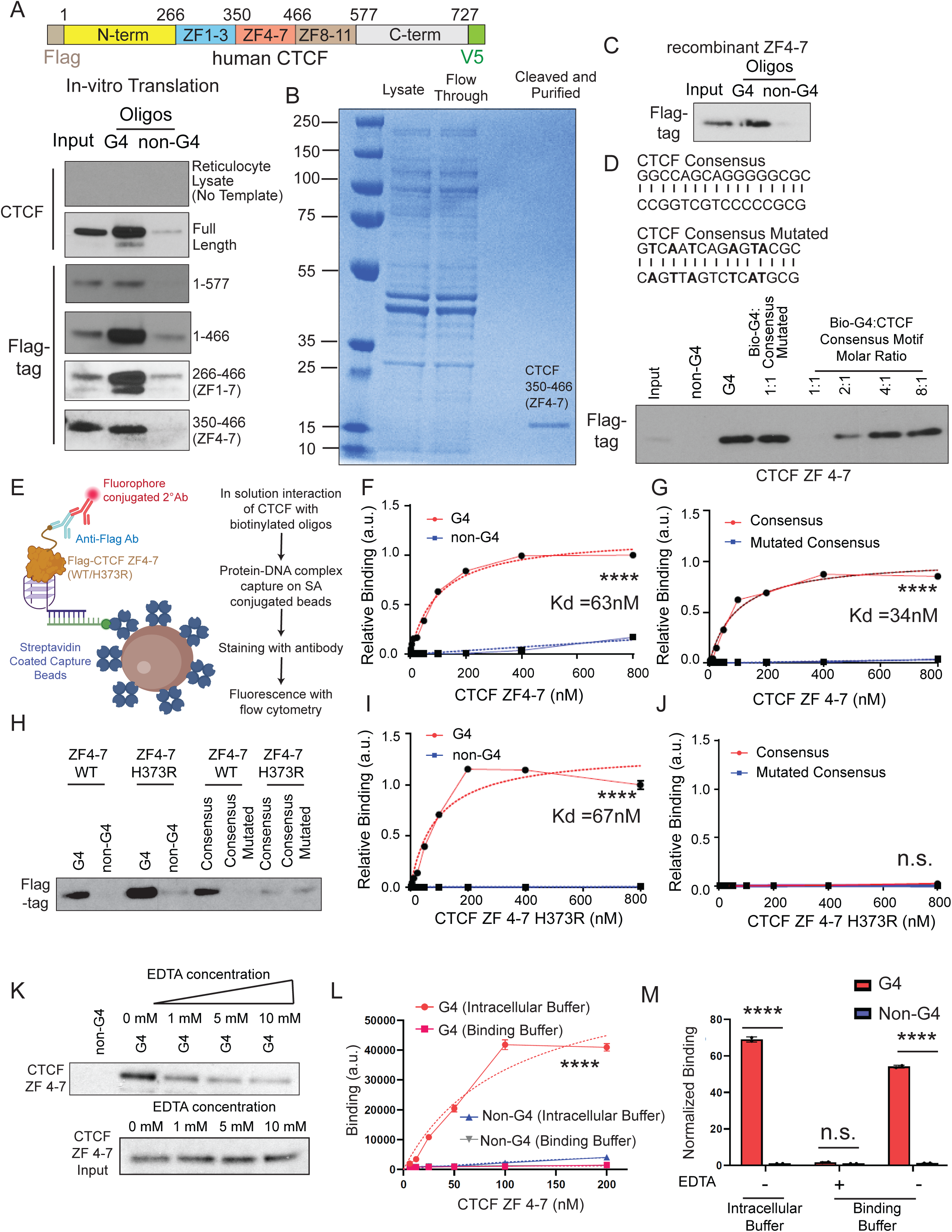
Biochemical validation of CTCF-G4 interaction A. Top: a schematic of the human CTCF with an N-terminal FLAG (beige) and C-terminal V5 tag (green) expressed using *in vitro* transcription and translation with rabbit reticulocyte lysates. Bottom: immunoblot showing the expression of *in vitro* transcribed and translated full length (FL) CTCF or mutants lacking specific domains. Pulldown was performed with biotinylated G4 oligos or non-G4 controls. Input without oligo pulldown was used as loading control, and reticulocyte lysate without template was used as negative control. B. Coomassie stained gel validating the expression and purification of recombinant CTCF fragments containing the central ZF4-7 fragment (CTCF 350-466) expressed in *E. coli*. C. Immunoblot showing the enrichment of recombinant CTCF ZF4-7 with G4 oligos, compared with non-G4 controls. D. Top: oligonucleotide sequence of CTCF consensus and mutated consensus motifs used in oligo pulldown. Each mutated base is bolded in the mutated consensus motif sequence. Bottom: Immunoblot showing the enrichment of recombinant CTCF ZF4-7 with biotinylated (*Kit2)* G4 compared to non-G4 control oligos. The biotinylated G4 oligo was mixed with varying molar ratios of dsDNA consensus motif or with an equimolar concentration of mutated consensus. E. Schematic of G4 binding assay performed using flow cytometry. Flag-tagged recombinant CTCF was mixed with different biotinylated oligos followed by capture on streptavidin (SA) magnetic beads. The specific interaction was quantified by on-bead fluorescence signal measured using an anti-Flag antibody. F. Binding curve for interaction between wild-type (WT) recombinant CTCF ZF4-7 and *Kit2* G4 (red, circle) or non-G4 (blue, square) oligos. The relative binding was calculated by normalizing the fluorescence signal of bead only negative control, stained with the anti-Flag antibody. X-axis denotes CTCF protein concentrations in nanomolar (nM), Y-axis represents arbitrary units (a.u.) of binding. Curved dashed lines represent the inferred non-linear fit for the data. G. Binding curve for interaction between WT recombinant CTCF ZF4-7 and consensus or mutated consensus oligos. X-axis denotes CTCF protein concentrations in nanomolar (nM), Y-axis represents arbitrary units (a.u.) of binding. Curved dashed lines represent the inferred non-linear fit for the data. H. Immunoblot showing the pull down of recombinant WT or H373R CTCF ZF4-7 with biotinylated G4, non-G4, consensus, or mutated consensus oligos. I. Binding curve showing the interaction between H373R recombinant CTCF ZF4-7 and *Kit2* G4 (red, circle) or non-G4 (blue, square) oligos. X-axis denotes CTCF protein concentrations in nanomolar (nM), Y-axis represents arbitrary units (a.u.) of binding. Curved dashed lines represent the inferred non-linear fit for the data. J. Binding curve showing the interaction between H373R recombinant CTCF ZF4-7 protein and consensus (red, circle) or mutated consensus (blue, square) oligos. X-axis denotes CTCF protein concentrations in nanomolar (nM), Y-axis represents arbitrary units (a.u.) of binding. Curved dashed lines represent the inferred non-linear fit for the data. K. Immunoblot showing the pull down of recombinant CTCF ZF4-7 with biotinylated G4 oligos in presence of varying concentrations of EDTA (0mM to 10mM). Input without oligo pulldown was used as loading control (bottom) and lysate from protein pulled down with non-G4 was used as negative control. L. Binding curve showing the interaction of recombinant CTCF ZF4-7 with *Kit2* G4 or non-G4 oligos in either intracellular buffer (orange circle and blue triangle, respectively) or binding buffer from Wulfridge et.al. (red square and grey triangle, respectively). X-axis denotes CTCF protein concentrations in nanomolar (nM), Y-axis represents arbitrary units (a.u.) of binding. Curved dashed lines represent the inferred non-linear fit for the data. ****p value ≤ 0.0001 calculated with 2-way ANOVA M. The effect of EDTA on binding of the recombinant CTCF ZF4-7 protein to *Kit2* G4 (red) or non-G4 (blue) oligos in either intracellular or binding buffer. The binding was normalized to bead only negative controls. Data points represent the mean and ± standard error from three independent experiments. **** p value ≤ 0.0001 calculated with student T-test. n. s. Not significant. Data points in figures F, G, I and J represent the mean ± standard deviations from three independent experiments. **** p value ≤ 0.0001 calculated with 2-way ANOVA.

Since CTCF ZFs 4-7 mediates interaction with both G4 and dsDNA consensus motifs, we compared these binding activities in a competition assay. For these experiments, we mixed the biotinylated G4 oligo (*Kit2*) with either the (non-biotinylated) CTCF consensus dsDNA motif or a mutated version of the dsDNA consensus at different molar ratios (**Fig.2D**). Following in-solution binding of CTCF ZF4-7 with different oligo mixtures we recovered G4 bound fractions through pulldown with streptavidin conjugated beads. While the mutated dsDNA consensus motif did not affect G4 bound CTCF fraction, an equimolar ratio of the consensus dsDNA motif completely abolished CTCF binding to G4 (**Fig.2D**). Notably, the G4 bound CTCF fraction was progressively recovered with 2-fold molar excess of G4 and showed binding at saturation levels when 4-fold molar excess of G4 oligo was used (**Fig.2D**). Based on these results, we estimate that the binding affinity of CTCF ZF4-7 for G4 is approximately 2-4-fold lower than that for its dsDNA consensus motif.

To measure the binding affinities of CTCF ZF4-7 for G4 and consensus DNA, we then developed a binding assay in which CTCF interaction with different DNA substrates was measured under native (intracellular) salt concentrations (**Fig.2E**). For this, CTCF was incubated with biotinylated DNA substrates in solution, followed by streptavidin capture and on-bead fluorescence measurement using particle-based flow cytometry (**Fig.2E**). It is important to note that unlike electromobility shift assays (EMSAs) or other similar assays, this method allows measurement of binding affinities under more relevant salt concentrations in the absence of EDTA, which is a potent inhibitor for DNA binding activity of zinc finger proteins^46^, such as CTCF. As expected, CTCF ZF4-7 domains exhibited robust binding to G4 DNA (*Kit2*) with a dissociation constant (K_d_) of ∼63nM, whereas non-G4 oligo did not show any appreciable binding (**Fig.2F**). In comparison, CTCF ZF4-7 bound more strongly to its dsDNA consensus motif with K_d_ of ∼34nM, which is comparable to previous estimates using fluorescence polarization (**Fig.2E**)^32^. Therefore, the binding affinity of CTCF ZF4-7 for its consensus DNA motif is approximately two-fold higher than that for G4 DNA, which is consistent with estimates from the competition assays (**Fig.2D**).

To further compare the CTCF-G4 interaction to CTCF binding to the consensus DNA motif, we generated a single point mutation of histidine 373 to arginine (H373R), the second histidine in C2H2 motif in ZF4, which is critical for recognition of CTCF consensus motif (**Sup.Fig.1C**). As expected, the CTCF ZF4-7 H373R exhibited major deficits in its ability to recognize the CTCF consensus motif, whereas in contrast, the H373R mutant showed no defects in G4 recognition (**Fig.2H-J**). Though the same region of CTCF provides specificity for consensus motif and G4 DNA, these results suggest that CTCF likely recognizes G4 and linear DNA substrates through distinct mechanisms. To ask whether the Zn finger structures of CTCF are required for G4 recognition, we performed binding measurements of CTCF to G4 DNA in the presence of varying concentrations of a Zn^2+^ chelator EDTA (**Fig.2K**) and observed a concentration-dependent decrease in CTCF-G4 interaction with increasing amounts of EDTA (**Fig.2K, Sup.Fig.1D**). These findings suggest that the ZF domain structure of CTCF is indeed important for G4 recognition.

A recent study from Wulfridge et.al. proposed that G4s facilitate enhanced binding of CTCF to its consensus motif, partly because they did not detect direct interaction of CTCF with G4s^20^. To address this, we compared CTCF-G4 interaction in the intracellular salt buffer used in our binding assays to the binding buffer used for EMSA performed in this study^20^ (**Fig. 2L-M**). Notably, the binding of CTCF ZF4-7 to G4s was greatly reduced in the EMSA binding buffer (from Wulfridge et.al.^20^), which contained EDTA, and removal of EDTA substantially recovered G4 binding (**Fig. 2L-M**). Based on these results we reason that the failure to detect CTCF-G4 interaction is likely due to suboptimal binding conditions used in previous studies^20,22^.

### CTCF ZF4 mutant (H373R) associates with G-rich genomic regions

Next, to investigate the G4-dependent functions of CTCF on a genome-wide level, we used a previously described mouse embryonic stem cell (mESC) system in which GFP and auxin inducible degron (AID) tags are knocked into the endogenous *Ctcf* locus (AID-CTCF)^35^. These AID-tagged CTCF mESCs also harbors a the *tir* transgene, which encodes the Tir E3 ubiquitin ligase that ubiquitinates and degrades the endogenous AID-CTCF in the presence of the plant hormone, indole 3-acetic acid (IAA or Auxin), within hours (**Fig.3A**)^35^. We transduced the AID-CTCF mESCs with lentiviral expression plasmids for FLAG– and V5-tagged full-length (FL) CTCF, FL H373R mutant CTCF, or empty vector (EV) control, carrying mCherry as a reporter. The use of this experimental design avoids the problem that long-term CTCF depletion is detrimental to most cell types and allows us to develop a system in which CTCF-depleted mESCs could be reconstituted lentivirally to express FL CTCF or H373R mutant proteins.

**Figure 3.**
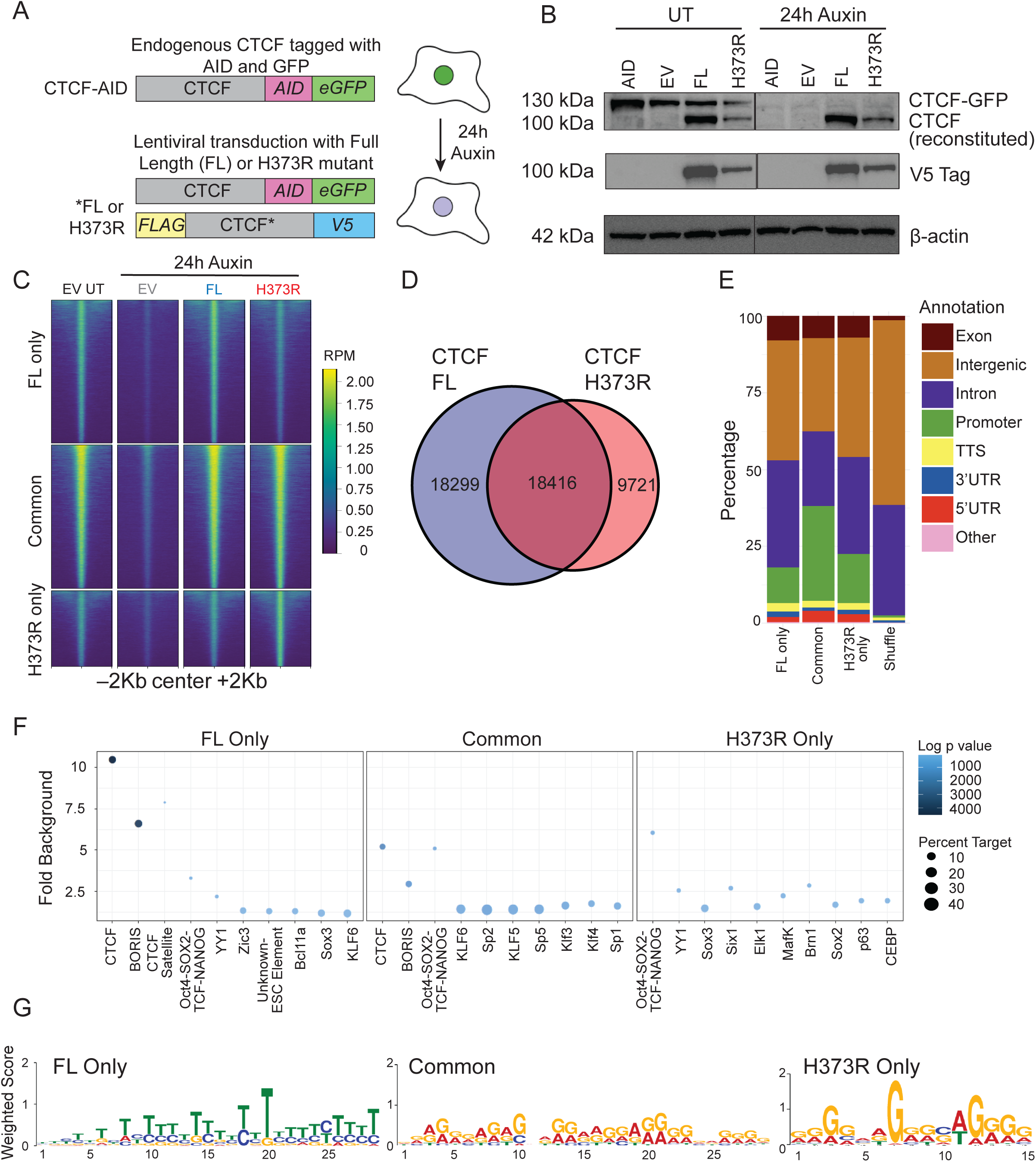
Genome-wide profiling of FL and H373R CTCF protein A. Schematic showing the endogenous CTCF tagged with Auxin Inducible Degron (AID) tag and GFP (top) and the exogenous Full Length (FL)* or H373R* CTCF construct tagged with FLAG (at N-terminus) and V5 (at C-terminus) (bottom). The cells were treated with auxin for 24 hours to monitor degradation of endogenous CTCF. B. Immunoblot showing the expression of GFP-tagged endogenous (130 kDa) or V5-expressing reconstituted (100 kDa) CTCF (top) with or without auxin treatment for 24 hours. The expression of reconstituted CTCF was validated with immunoblotting using anti-V5 antibody (middle). AID represents the original mESCs without auxin treatment. EV, FL and H373R are cells transduced with empty vector, full length and H373R CTCF, respectively. Beta-actin expression is used as loading control (bottom). Heatmaps showing CUT-and-Tag signal intensity of CTCF binding across ±2 kb regions centered on peaks with or without auxin treatment (24 hours) in EV, FL and H373R expressing cells. Peaks were classified into FL-only (top), Common (middle), and H373R-only (bottom) regions. CUT– and-Tag signal intensity, normalized to Reads Per Million (RPM), is depicted by the color scale. CTCF CUT-and-Tag was performed as three independent replicates across different experimental conditions. The heatmaps are merged/averaged signals from three replicates. C. Venn diagram showing the number of CTCF peaks identified using CUT-and-Tag in auxin-treated mESCs transduced with either FL (blue) or H373R (red) CTCF. D. Genomic annotations of CTCF peaks categorized as FL only, Common, and H373R only, along with genomic shuffle as control regions. E. Motif enrichment analysis of CTCF peaks categorized as FL-only (left), Common (middle), and H373R-only (right) regions. Y-axis represents the fold enrichment of motifs compared to the genomic background; X-axis denotes enriched motifs. Circle size indicates the percentage of peaks containing each motif, and the color intensity corresponds to the log-transformed p-value for motif enrichment performed using HOMER. F. Top sequence motifs identified using *de novo* motif discovery in FL-only (left), Common (middle), and H373R-only (right) CTCF peaks. The Y-axis represents the weighted score, indicating the contribution of each nucleotide within the motif.

We sort-purified transduced cells based on mCherry reporter (**Sup.Fig.2A**, *left*) and confirmed degradation of the 130 kDa CTCF-AID-GFP fusion protein expressed from the endogenous *Ctcf* locus was degraded at 8 or 24 hours of auxin treatment, while the reconstituted FL CTCF and H373R mutant proteins (detected by immunoblotting with anti-V5) were not degraded (**Fig. 3B, Sup.Fig.2B**). Since our biochemical studies were performed with CTCF ZF4-7 fragments, using in vitro translation assays, we confirmed that both FL and the H373R mutant CTCF binds G4 in the full-length context (**Sup.Fig.2C**, *top panel*). Moreover, as with the ZF4-7 fragment, the full length CTCF with H373R mutation also showed reduced binding to the CTCF consensus DNA motif (**Sup.Fig.2C**, *bottom panel*).

To determine whether the expression of the H373R mutant was deleterious to cells, we characterized the AID-CTCF mESCs post-auxin treatment. There were no significant differences in cell numbers (**Sup.Fig.2E**, *left panels*) of cells expressing empty vector or WT and mutant CTCF, or spontaneous differentiation to SSEA1-negative fibroblasts (**Sup.Fig.2F-G**) at 24 hours. However, by 72 hours of auxin treatment, CTCF depletion upon auxin treatment led to a significant reduction in cell numbers of the parental CTCF-AID-GFP-expressing cells and cells transduced with empty vector, but this loss of viability was rescued to similar extents by reconstitution with FL CTCF or H373R mutant (**Sup.Fig.2E**, *right panels*). Notably, the FL mutant CTCF H373R was expressed at lower levels than FL wildtype CTCF (**Fig. 3B**, *top and middle panels*; **Sup.** Fig 2C), even in cells sorted for equivalent levels of mCherry expression (**Sup.Fig.2A**, *right*) suggesting that the mutant CTCF protein may be less stable compared to the wildtype FL CTCF. Together these experiments indicate that the H373R mutant is capable of supporting the survival of CTCF-AID-GFP-depleted cells as efficiently as wildtype CTCF.

We then profiled the genomic distribution of CTCF using CUT-and-Tag (<u>C</u>leavage <u>U</u>nder <u>T</u>argets and <u>Tag</u>mentation) in AID-CTCF mESCs not treated with auxin or reconstituted with EV, FL CTCF or H373R mutant and treated with auxin for 24 hours. We noted a clear loss of CTCF binding in CTCF-depleted (EV) mESCs treated with auxin, and CTCF binding was recovered in cells expressing FL or H373R mutant (**Fig.3C**). The majority of peaks were shared between FL and H373R mutant (n=18,416) but the H373R mutant showed fewer binding peaks (n=28,137) compared to FL CTCF (n=36,715) (**Fig.3C**). We categorized all CTCF peaks from cells expressing FL and H373R mutant into sites preferentially bound by FL CTCF (FL only, n=18,299), H373R CTCF (H373R only, n=9,721) and those that were similarly enriched for FL and H373R binding (common, n=18,416) (**Fig.3D**). Genomic annotations of these regions showed increased representation of promoters and 5’ untranslated regions (5’UTRs) within common sites bound by both FL and H373R CTCF in comparison to FL only, H373R only or randomly shuffled genomic regions (shuffle) (**Fig.3E**).

We performed motif enrichment analysis on FL only, common and H373R only regions, and found the canonical CTCF motif as the most enriched motif in FL only and common regions (**Fig.3F**). The number of peaks with the canonical CTCF motif were reduced by two-fold in common regions as compared with FL only regions. On the other hand, consistent with the diminished ability of the H373R mutant to recognize the CTCF consensus motif, the CTCF motif was not among the top enriched motifs in H373R peaks (**Fig.3F**). Besides CTCF motifs, the FL only, common and H373R only peaks displayed enrichment of OCT4-SOX2-TCF-NANOG motifs, which is in line with previous studies demonstrating cooperativity of these transcription factors with CTCF (**Fig.3F**)^47^. We also performed *de novo* motif discovery analysis and observed enhanced GC content (**Sup.Fig.3A**) as well as enrichment of repetitive G4-like motifs (**Fig.3G**), in common and H373R only peaks. Corroborating the biochemical studies, these analyses highlight the reduced affinity of the H373R CTCF mutant to consensus CTCF motifs and reveal a predisposition towards G-rich genomic sequences.

### Common sites bound by FL and ZF4 mutant CTCF are enriched for G4s

Having observed an enrichment of G-rich sequences in regions bound by both FL and H373R CTCF, we then mapped G4s. G4-specific antibodies, 1H6 and BG4, have been the mainstays for genomic G4 profiling^44,48^. Though these antibodies show good affinity for G4 DNA, they have also been reported to cross-react with cytosine-rich and thymine-rich single-stranded (ss) DNA^49–51^. In our recent work^15^, we used a small molecule, N-methyl mesoporphyrin IX (NMM) as a detection probe for G4s in cells, which binds G4 in 1:1 stoichiometry through a well-defined π-stacking mechanism (**Sup.Fig.3B**)^52,53^. Importantly, NMM is highly selective for G4s, particularly G4s of parallel topologies, which motivated us to further develop it as a G4-specific pulldown probe (**Fig.4A**). Using propylphosphonic anhydride (T3P) as a coupling agent, we conjugated pegylated biotin amine with the carboxyl groups on NMM, which do not participate in G4 recognition by NMM (**Fig.4A**). We confirmed the generation of NMM-biotin complex (NMM-Bio) with mass spectrometry and purified NMM-Bio with chromatography. Upon titration, ∼10µM NMM-Bio efficiently pulled down G4 DNA and showed no detectable affinity for non-G4 DNA (**Fig.4B, Sup.Fig.3C**).

**Figure 4.**
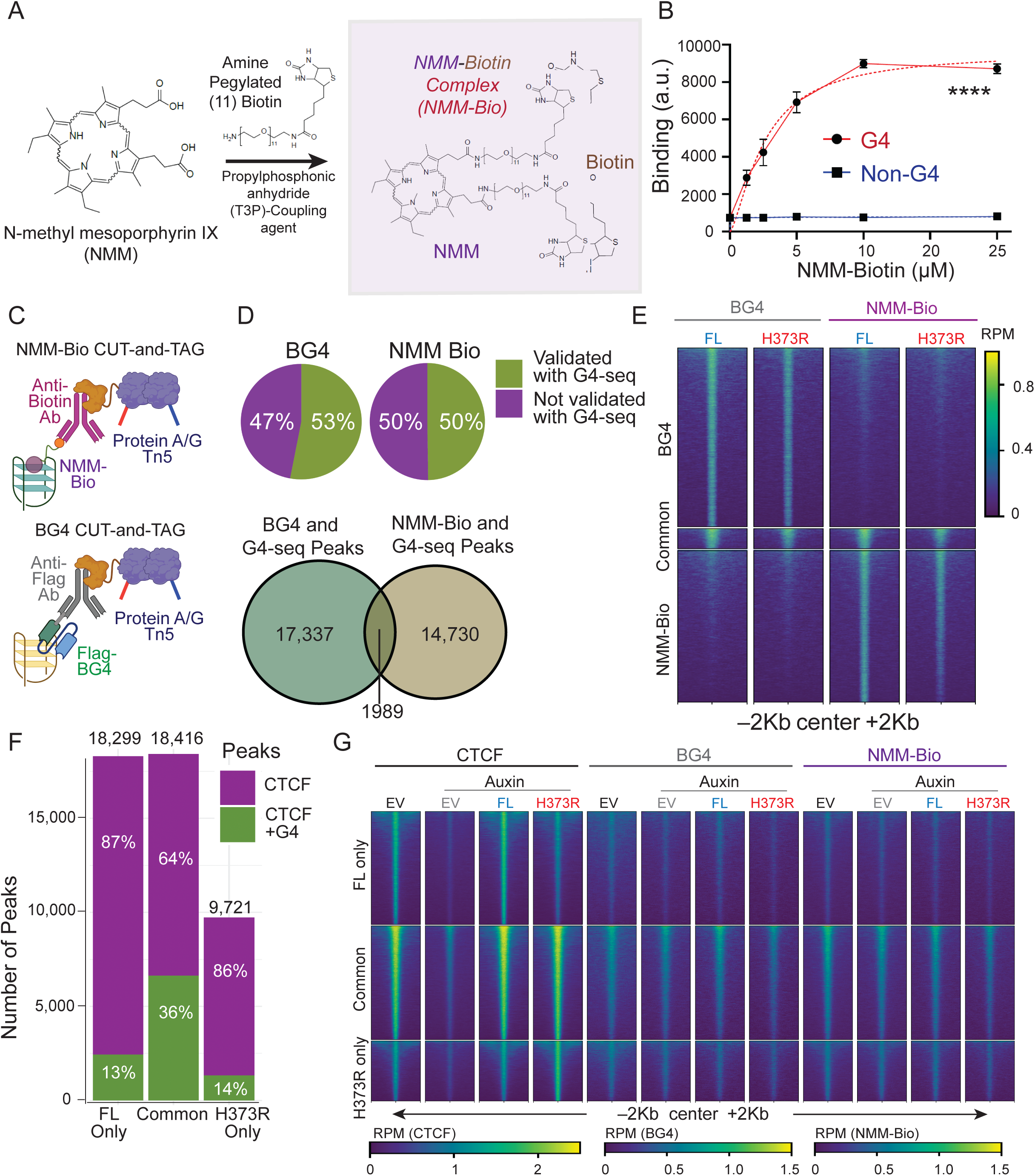
Genome-wide G4 profiling and its association to CTCF binding A. Schematic for the generation of N-methyl mesoporphyrin IX (NMM) conjugated with biotin (NMM-Bio). B. Binding curve of NMM-Bio to G4 (red) and non-G4 (blue) oligos. X-axis marks different NMM-Bio concentrations used, and Y-axis represents binding signal as arbitrary units (a.u.). Data points are mean ± standard error from four independent experiments. **** p value ≤ 0.0001 calculated with 2-way ANOVA. C. Schematic representation of CUT-and-TAG for NMM-Bio and BG4 based G4 profiling. NMM-Bio was combined with anti-biotin antibody, while flag-tagged BG4 single chain variable fragment antibody was combined with anti-Flag antibody. In both methods, tagmentation was performed using Tn5 transposase fused with protein A/G. D. Overlap of BG4 and NMM-Bio peaks with previously published G4-seq peaks identified in mouse genomic DNA. Top: pie charts show the percentage of BG4 and NMM-Bio peaks with or without G4-seq overlap. Bottom: Venn diagram highlights shared and unique peaks between BG4 and NMM-Bio methods at regions that are common with G4-seq regions in both conditions. E. Heatmaps showing signal intensity for BG4 and NMM-bio peaks ±2 kb from the center. Peaks were classified into unique and shared regions identified by NMM-Bio and BG4 methods in mESCs expressing FL and H373R CTCF upon 24 hours of auxin treatment. CUT-and-Tag signal intensity normalized to Reads Per Million (RPM), is depicted by the color scale. F. Bar plot quantifying the overlap of G4 peaks (identified by either BG4 or NMM-Bio method) with CTCF peaks categorized into FL-only, Common, and H373R-only regions. G. Heatmaps showing signal intensity (RPM) of CTCF, BG4, and NMM-Bio signals across FL-only, Common, and H373R-only CTCF peaks. Regions were classified into FL-only, Common, and H373R-only peaks in mESCs expressing EV, FL and H373R CTCF with or without (EV control) auxin treatment. BG4 and NMM-Bio CUT-and-Tag were performed as three independent replicates across different experimental conditions. Unconjugated NMM (NMM) was used as negative control, and the peaks were called using NMM signal as background. The heatmaps are merged signals from three replicates.

We then adapted NMM-Bio in CUT-and-Tag workflow and profiled genomic G4s using both BG4 and NMM-Bio CUT-and-Tag and used unconjugated NMM combined with anti-biotin antibody as control (**Fig.4C**). We first asked whether the peaks identified by BG4 and NMM-Bio CUT-and-Tag in FL or H373R CTCF expressing mESCs overlapped with G4s identified using G4-seq in non-chromatinized mouse genomic DNA^7,54^. Genomic peaks identified by BG4 and NMM-Bio showed 53% and 50% overlap with regions marked by G4-seq, respectively (**Fig.4D**). The regions identified by G4-seq showed only a minor overlap (∼10%) with G4 peaks identified by BG4 and NMM-Bio CUT-and-Tag, suggesting that BG4 and NMM-Bio identify distinct sets of genomic G4s in cells (**Fig.4D-E**). Comparison of BG4 and NMM-Bio CUT-and-Tag peaks overlapping G4-seq peaks showed a similar distribution of predicted G4s of distinct subtypes across methods (**Sup.Fig.4A**). Considering that BG4 and NMM-Bio methods reveal unique G4 regions, we decided to focus on the union of regions identified by either of these methods (i.e. 1989 peaks) for subsequent analysis.

To examine the association between genomic CTCF binding sites with G4 forming regions, we overlapped FL only, common, and H373R only CTCF sites with G4 regions identified using BG4 or NMM-Bio. In concordance with the G4 binding ability of FL and H373R, we observed most overlap of G4 peaks with the common CTCF peaks (36%, n=6,630), that were independently bound by both FL and H373R CTCF (**Fig.4F-G**). Even though the H373R mutant is expressed at a lower level and shows fewer statistically defined peaks, the proportion of H373R peaks overlapping G4 peaks was slightly higher than FL CTCF peaks (28% of H373R peaks vs 25% of FL peaks) (**Sup.Fig.4B**). We also analyzed ChIP-Seq data from a published study^39^, where several ZFs of CTCF were individually mutated by histidine (H) to arginine (R) substitutions and overexpressed in murine B cells with a BioID tag (**Sup.Fig.4C**). We found enrichment of a strictly defined G_3_N_1-7_G_3_N_1-7_G_3_N_1-7_G_3_ (3G-G4 motif) at CTCF (wildtype) binding sites (10% of peaks with 3G-G4 motif). Interestingly, the enrichment of 3G-G4 motif was further increased with mutations in ZFs 4 to 8 (from 10% to ∼30-40%), suggesting a shift in CTCF binding to regions with a higher propensity to form G4s. Additionally, we found an increase in total number of CTCF binding sites (peaks) when G4s were stabilized in CH12 B cells using small molecule ligands pyridostatin (10µM PDS, 17,331 peaks) and 3,11-Difluoro-6,8,13-trimethyl-8H-quino[4,3,2,kl] acridinium methosulfate (2µM RHPS4, 17,458 peaks) when compared to DMSO (10,053 peaks) (**Sup.** Fig. 4D). Overall, these experiments reveal clear genome-wide associations of CTCF binding with G4 structures.

### CTCF directly interacts with genomic G4 structures

To further dissect the CTCF-G4 interaction on a genomic level, we categorized CTCF peaks commonly bound by both FL and H373R CTCF into those with or without a canonical CTCF motif (Homer). As expected, from the diminished ability of H373R mutant to bind CTCF consensus DNA, regions lacking the canonical CTCF motif showed higher enrichment of G4 peaks (**Sup.Fig.4E,4F**). Within common CTCF peaks (bound by FL and H373R CTCF) that overlapped G4 peaks, we calculated the distance between the center of CTCF and G4 peaks (**Fig.5A**). Notably, a small number of regions with overlapping CTCF and G4 peaks that harbored canonical CTCF motifs (n=1556), exhibited significantly higher median distance between the centers of CTCF and G4 peaks, compared to regions which lacked a canonical CTCF motif (n=5467) (**Fig.5A**). Though these results infer that G4 structures and CTCF binding tightly coincide at sites lacking a defined canonical CTCF motif, the enrichment-based approaches such as CUT-and-Tag are not ideal for interrogating the CTCF-G4 interaction at base-resolution. To this end, we classified overlapping CTCF and G4 peaks into those that harbor only a single G_3_N_1-7_G_3_N_1-7_G_3_N_1-7_G_3_ (3G-G4) motif in the with or without a canonical CTCF motif. Interestingly, in regions (n=279) which carried single CTCF and 3G-G4 motifs, the average distance between the two motifs was >300 bp (**Fig. 5B**). Moreover, the average distance of the 3G-G4 motif to the nearest CTCF motif was ∼5kb at overlapping CTCF and G4 peaks that otherwise lacked a canonical CTCF motif (**Fig. 5C**). Together, these analyses highlight two important points: 1) at several places in the genome CTCF binding directly coincides with G4 sequences, and 2) even at CTCF binding sites where G4 and CTCF motifs co-exist, on average these motifs are separated by a few hundred base pairs.

**Figure 5.**
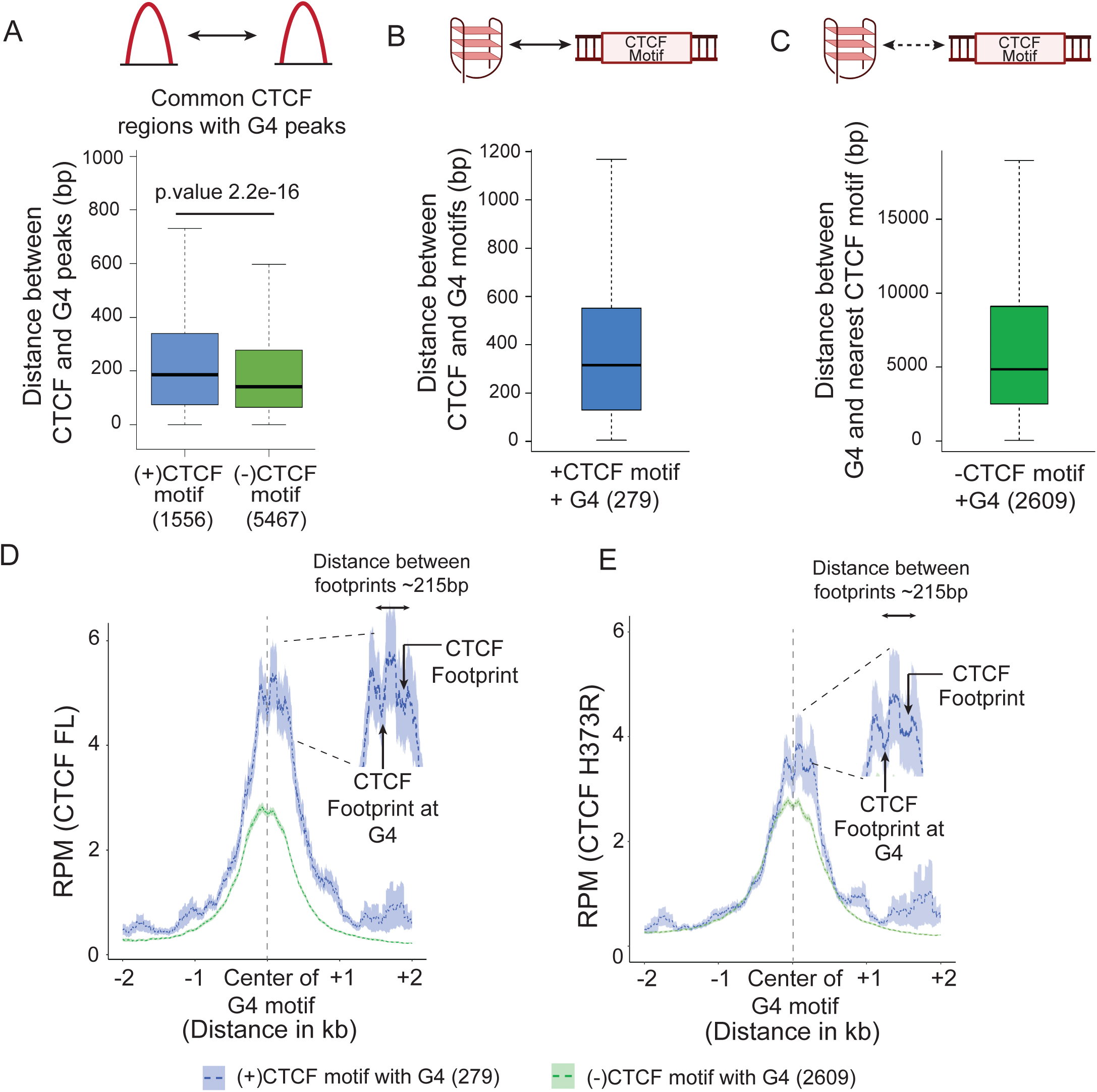
CTCF directly associated with genomic G4s A. Box and whisker plot quantifying the distance between the centers of the overlapping CTCF and G4 peaks. The regions were further separated based on the presence (+CTCF motif) or absence (-CTCF motif) of a canonical CTCF motif within the peak. The statistical significance was calculated using Wilcoxon signed-rank test. B. Box and whisker plot quantifying the distance between the center of the CTCF motifs (+CTCF motif) and the 3G-G4 motif within common (FL and H373R enriched) CTCF peaks that overlap a G4 peak. C. Box and whisker plot quantifying the distance between the 3G-G4 motif to the nearest CTCF motif in common (FL and H373R) CTCF peaks that overlap a G4 peak. D. Histograms displaying the signal from DNA footprinting analysis at overlapping CTCF and G4 peaks separated into regions with (blue) or without (green) a canonical CTCF motif. The FL CTCF footprints are centred at the 3G-G4 motif within G4 peaks. The zoomed in area is the center of G4 peak with canonical CTCF motif. The shaded area represents standard deviation. E. Histograms displaying the signal from DNA footprinting analysis at overlapping CTCF and G4 peaks separated into regions with (blue) or without (green) a canonical CTCF motif. The H373R CTCF footprints are centred at the 3G-G4 motif within G4 peaks. The zoomed in area is the center of G4 peak with canonical CTCF motif. The shaded area represents standard deviation.

To assess this more carefully, we examined the footprints of FL and H373R CTCF binding at overlapping CTCF (common regions) and G4 peaks. For these analyses we plotted the signal for CTCF binding relative to the center of the 3G-G4 motif in regions categorized based on the presence or absence of a canonical CTCF motif. Remarkably, the binding footprint of CTCF precisely matched the center of the G4 motif, specially at sites lacking a canonical CTCF motif (**Fig.5D-E**). Furthermore, at regions that harbored canonical CTCF motifs, two distinct CTCF footprints were evident (**Fig.5D-E**). One of these CTCF footprints directly overlaid the center of G4 motifs and the other was located ∼215 bp (mean distance) downstream to the G4 motif (**Fig.5D-E**). The CTCF footprint at the center of G4 motifs was distinct in its appearance (narrower) when compared to the downstream CTCF footprint, which we presume to be mediated by the canonical CTCF motif (**Fig.5D-E**). Both FL CTCF and H373R mutant exhibited these unique binding features (**Fig.5D-E**). However, compared to FL CTCF, the signal for H373R binding was visibly reduced in regions harboring the canonical CTCF motif and enhanced at regions that lacked the canonical CTCF motif (**Fig.5D-E**). From these results we propose that CTCF directly binds G4 structures at least at a subset of genomic sites.

### CTCF-G4 interaction mediates chromatin looping and is associated with changes in gene expression

To elucidate the role of CTCF-G4 interaction in regulation of gene expression, we performed RNA-seq in cells expressing empty vector (EV) control with or without auxin and mESCs reconstituted with FL or H373R mutant 24 hours post auxin mediated CTCF depletion. Principal component analysis showed clustering of all replicates, with PC1 accounting for 73% of the variance and segregating the samples based on auxin treatment (**Fig.6A**). Notably, PC2 further stratified samples based on CTCF expression, highlighting the distinct transcriptional responses upon CTCF reconstitution. Differential gene expression analysis in EV expressing samples treated with or without auxin revealed 978 differentially expressed genes (DEGs; false discovery rate ≤0.05, log2 fold change > ±0.58) with 520 up– and 458 down-regulated genes (**Sup.Fig.5A**). The comparison between FL and EV expressing cells treated with auxin identified 799 DEGs (361 up and 438 down), whereas H373R versus EV expressing cells showed 431 DEGs (189 up and 242 down) upon auxin treatment (**Fig.6B**). Of these DEGs, 233 DEGs (97 up and 136 down) were shared in pairwise comparisons of FL and H373R to EV controls (**Fig.6C-E**). We reasoned that given the common binding activity of FL and H373R CTCF towards G4s, these 233 DEGs might be enriched for G4 signal. Indeed, the 233 DEGs exhibited enrichment of CTCF, BG4, and NMM-Bio signals at their transcription start sites (TSSs) (**Fig.6F**). 38% of these genes (89/233) also harbored statistically defined CTCF and G4 peaks in their vicinity (± 10kb of the gene), suggesting that these are likely G4-regulated genes. Gene ontology analysis of the shared DEGs between FL and H373R showed that upregulated genes were enriched for pathways related to embryonic development, cell fate commitment and vitamin metabolism, whereas downregulated genes were associated with developmental programs linked to organogenesis (**Fig.6G**). These data suggest that both FL and H373R CTCF are essential for maintaining commitment to an embryonic stem cell program and repressing sporadic activation of organ/lineage specific developmental pathways.

**Figure 6.**
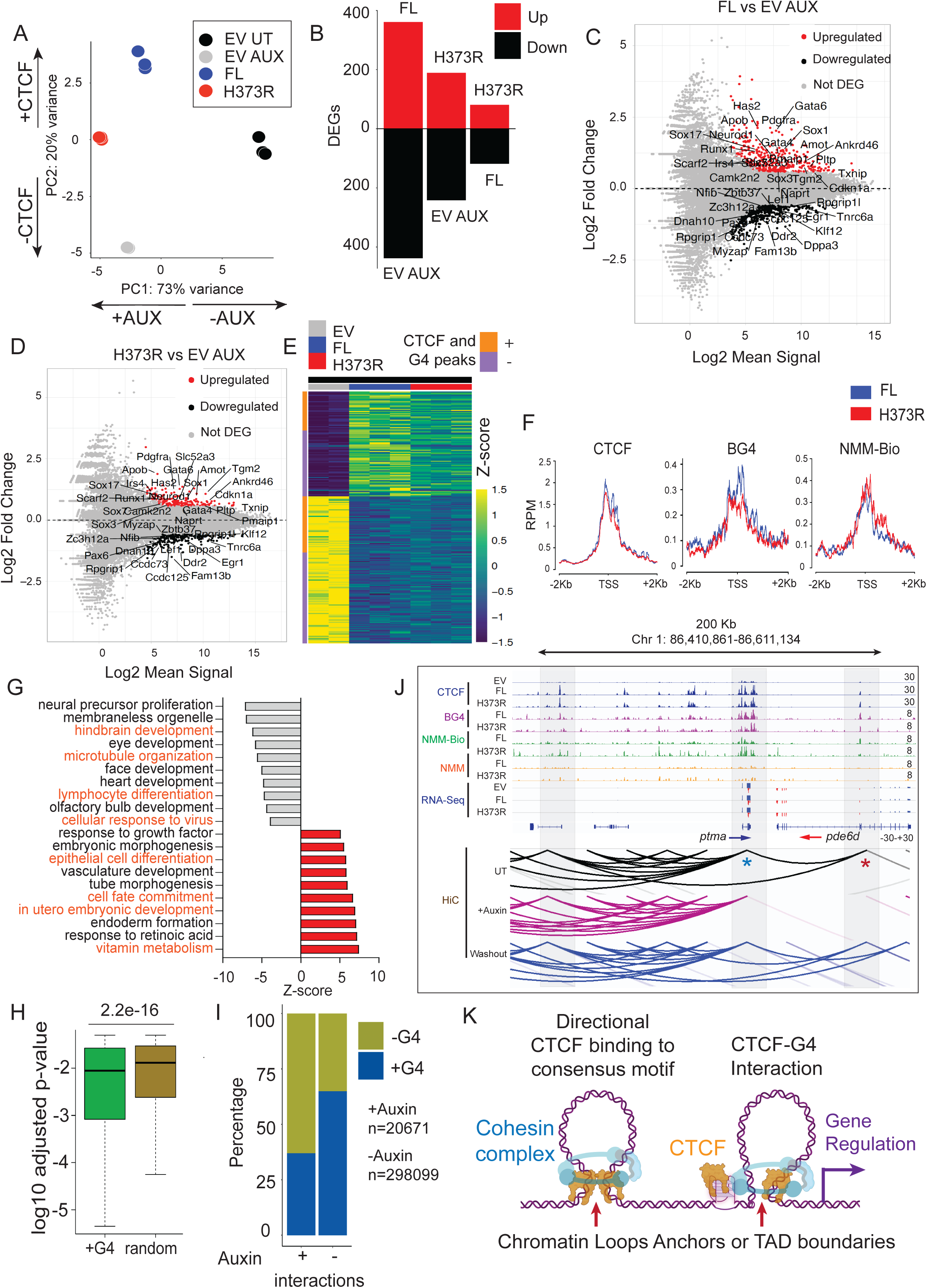
CTCF-G4 interactions mediate specific gene expression changes A. Principal component analysis of RNA-seq data of mESCs with EV untreated (UT) (black), EV auxin (AUX) (gray) and FL auxin (blue) and H373R auxin (red). Percent variance for principal components 1 (PC1) and 2 (PC2) are shown on X and Y-axis respectively. Arrows in each axis represent the features explaining the variance: with or without auxin (PC1) and presence or absence of CTCF (PC2). B. Barplot showing the numbers of differentially expressed genes (DEGs) in pairwise comparisons between full length (FL) vs Empty vector (EV)-AUX, H373R vs EV-AUX and H373R vs FL. Genes are separated into up (red) and downregulated (black) C. MA plots showing the DEGs in FL CTCF vs EV-AUX. Red and black points are upregulated and downregulated genes respectively. X-Axis represents Log2 base mean signal and Y-axis denotes Log2 fold change in expression. D. MA plots showing the DEGs in H373R CTCF vs EV-AUX. Red and black points are upregulated and downregulated genes respectively. X-Axis represents Log2 base mean signal and Y-axis denotes Log2 fold change in expression. E. Heatmap showing the Z-score from the RPKM expression values of 233 common DEGs between FL vs EV-AUX and H373R vs EV-AUX comparisons. F. Profile plot showing the CTCF, BG4 and NMM-Bio signal as Reads Per Million (RPM) at the transcription start sites (TSS) ±2 kb within the 233 common DEGs in FL vs EV-AUX and H373R vs EV-AUX. G. Downregulated (gray) and upregulated (red) pathways in the 233 common DEGS in FL vs EV-AUX and H373R vs EV-AUX. X-axis represents Z-score. Pathways marked in orange are enriched for genes that show overlap with both CTCF and G4 peaks. H. Box and whisker plot comparing log10 p-values from Hi-C interactions (untreated condition) that overlap G4 peaks (green; G4) or genomic shuffle regions (random; olive) as controls. The statistical significance was calculated using Wilcoxon signed rank test. I. Barplot quantifying Hi-C interactions with or without auxin treatment (for 48 hours) in AID-tagged mESCs. The Y-axis represents the percent of interactions with (blue) or without (olive) an overlapping G4 peaks identified using NMM-Bio or BG4 in mESCs reconstituted with FL or H373R CTCF treated with auxin. The unique interactions that remained post-auxin treatment (n=20,671) were compared to unique interactions in untreated conditions (n=298, 099). J. Representative genome browser view of CTCF (blue), BG4 (purple), NMM-Bio (Green), NMM (Orange) and RNA-seq (blue/red) at *Ptma* and *Pde6d* gene loci. The expression of *Pde6d* gene is positively regulated by FL CTCF and H373R mutant. Shaded areas define regions of interest with CTCF and G4 peaks. Bottom part is HiC contact maps from Nora et.al.^35^ in the parent mESCs without auxin (untreated, UT), with auxin treatment for 2 days (+Auxin) and 2 days following auxin washout (Washout). The red and blue asterisks highlight the interaction anchor points that are lost or retained upon auxin treatment. The same parent mESCs were used in our studies. K. A proposed model depicting the role of CTCF-G4 interaction in mediating long-range chromatin looping to define stable chromatin loops or boundaries of topologically associating domains (TADs).

We also performed clustering of transcriptomics data with time-course sequencing (TC-seq) analysis to study the relationship between CTCF binding, G4 formation and dynamic changes in gene expression patterns in different experimental conditions. TC-seq generated 9 unique clusters, C1 through C9 (**Sup.Fig.5B-C**). We narrowed these clusters based on expression changes in response to CTCF depletion with auxin and further sorted the clusters considering similar effects of FL and H373R reconstitution on restoration of gene expression patterns. Among the interesting clusters, cluster C1 was upregulated by CTCF, whereas cluster C2 was seemingly repressed by CTCF (**Sup.Fig.5B-C**). Moreover, clusters C1 and C2 were enriched in pathways related to immune response and neurogenesis, respectively. Notably, in clusters C1 and C2, reconstitution with FL CTCF restored expression profiles resembling the untreated (UT) EV controls (**Sup.Fig.5B-C**). Reconstitution with H373R mutant also restored the gene expression patterns, albeit the effects were less prominent, which could be due to the lower expression levels of H373R compared to FL (**Sup.Fig.5B-C; Fig.3B**). Nonetheless, the CTCF, BG4, and NMM-Bio signals within ±200bp of the TSS of genes in clusters C1 and C2 also showed an observable enrichment when compared to the non-auxin responsive cluster C4 as control (**Sup.Fig.5D**).

Finally, to link the changes in gene expression to CTCF-mediated long-range chromatin looping, we analyzed a previously published high throughput chromosome conformation capture (Hi-C) dataset which was generated in the AID-tagged CTCF mESCs, the same parent line used in our studies. We generated chromatin contact maps reporting significant interactions with (auxin) or without (untreated, UT) auxin treatment for 48 hours, as well as 48 hours post-auxin washout (washout). Using FitHiC, an algorithm for identifying significant chromatin interactions, we identified a total of 318,770 chromatin interactions in untreated cells at 20kb resolution, and upon auxin treatment a large proportion of these interactions were lost (n=66,073 remaining post-auxin) (**Sup.Fig.5E**). Importantly, the majority of interactions were recovered post-auxin washout (n=328,783) (**Sup.Fig.5E**). Using these interactions, we first asked if the presence of G4s affects the strength of chromatin contacts (**Fig.6H**). Indeed, consistent with previous reports^19,20^, interactions that overlapped with a G4 peak showed a significantly lower adjusted p-value, when compared to randomly surveyed interactions without G4s (**Fig.6H**). These data suggest that chromatin loops that are associated with a G4 may be functionally distinct from the ones that do not carry a G4. To assess this further, we compared the G4 distribution in all reported Hi-C interactions in untreated mESCs and found that ∼35% interactions were associated with G4 peaks (**Fig. 6I, (-)auxin**). Remarkably, the G4 peaks were even more enriched (∼66%) at chromatin loops that persisted upon CTCF depletion with auxin treatment (**Fig.6I, (+)auxin**), we refer to these as “persistent loops”. To rule out the possibility that CTCF binding is retained at these sites (persistent loops) even upon auxin treatment, we plotted the CTCF signal in (CTCF) peaks that overlapped Hi-C interaction windows in untreated conditions or those that persisted upon auxin treatment in our datasets. As expected, the CTCF signal was similarly depleted upon auxin treatment (EV AUX) in CTCF peaks associated with chromatin loops from untreated or auxin conditions (**Sup.Fig.5F**; grey lines). We noted several examples of these “persistent loops” in the genome, including a region around the *Pde6d* and *Cldn3* gene loci, where CTCF binding overlapped G4 containing regions and while some chromatin contact loop anchors were lost upon auxin treatment, others remained intact (**Fig.6J, Sup.Fig.6A**). Importantly, many of these sites were associated with gene expression changes that were linked to FL and H373R CTCF binding. 197/233 DEGs co-regulated by CTCF FL and H373R mutant were associated with Hi-C interactions with CTCF and G4 peaks, and intriguingly, 107/233 DEGs were associated with persistent loops, displaying a highly significant correlation (<0.00001, Chi-square test) (**Sup.Fig.5G**). In contrast to the loops that persisted upon auxin treatment, several other chromatin contacts were totally obliterated upon auxin treatment at certain genomic loci, such as the *Cox15* gene locus, that also showed association with CTCF binding and G4s at a distant regulatory region (**Sup.Fig.6B**). In conclusion, our findings suggest that G4s, in conjunction with CTCF binding, play a critical role in regulation of gene expression possibly by influencing long-range chromatin interactions.

## Discussion

Our study expands the understanding of DNA G4 structures as critical regulators of genomic function, with particular focus on their direct interaction with the genomic architectural protein, CTCF. By integrating proteomic screens and biochemical assays, we provide strong evidence for the specific recognition of G4s by the central ZF domains of CTCF. Through genome-wide analyses, we suggest that CTCF recruitment may be directly mediated by G4s at 15-20% of genome-wide CTCF binding sites; this is in addition to the ∼80 to 85% of CTCF binding sites that contain consensus CTCF motifs^38,39^ With these findings, we propose a model in which CTCF-G4 interaction cooperates with directional CTCF binding to its consensus motif for establishing chromatin loops and TAD boundaries to regulate gene expression and possibly other genomic processes (**Fig.6K**). This hypothesis is supported by our finding that the anchor points of chromatin loops associated with G4s appear to be highly persistent and are less prone to CTCF depletion. This suggests that G4 structures and the associated DNA conformation by themselves may function as efficient loop anchors, even when CTCF is depleted. Alternatively, the G4 structures may recruit other proteins that may continue to serve a barrier function, even in the absence of CTCF. Taken together, our studies highlight the architectural function of G4 structures in the genome and provide fascinating insights into the paradigm of 3D genome organization.

The N-terminal region of CTCF is shown to specifically interact with the STAG2 (SA2) subunit of cohesin ring complex to impede loop extrusion^55,56^. It is important to note, that the cohesin ring complex is estimated to have a diameter of ∼40 nm^57^, whereas the approximate size of CTCF is 3-5 nm. Therefore, from a physical perspective, how CTCF even when bound to its DNA motif in the right orientation, can serve as an effective roadblock for cohesin loop extrusion which occurs at an estimated 1kb/second^37^, is unclear. In this regard, G4 structures and the looped out DNA strand with associated DNA:RNA hybrids could function as larger physical impediments for movement of cohesin rings. Moreover, a recent report has accentuated the importance of DNA tension as a key parameter for the barrier activity of CTCF^37^. By partially unwinding DNA strands and creating torsional stress, G4s can modulate DNA tension and may contribute to the barrier activity of CTCF by influencing this parameter^58^. It is also important to point out that besides CTCF, another genomic architectural protein, Yin-Yang1 (YY1) has been shown to bind G4 structures^21^. When considered together, these studies suggest that G4s have perhaps co-evolved for for interactions with DNA-binding proteins to regulate long-range DNA looping.

The interaction of CTCF with G4 structures has been controversial. Our work indicates different assay conditions as the possible source of discrepancies reported in previous studies^20,22,24^. We show that similar to other zinc finger containing proteins^46^, the binding of CTCF to G4s is sensitive to EDTA. We therefore reason that the inability to detect CTCF-G4 interaction in previous studies could be attributed to the presence of EDTA in binding and assay buffers^20,22^. Studies by Wulfridge et.al. suggested that G4 structures facilitate stronger binding of CTCF to weaker genomic motifs^20^. While this could be relevant for CTCF binding at certain genomic sites, this model requires precise positioning of G4s relative to the CTCF DNA motifs. Further, CTCF binding is disrupted by DNA methylation within the consensus DNA motifs^32,59^, and G4 structures have been linked to inhibition of DNA methylation through direct binding and inhibition of DNMT1^14^. Therefore, as an alternative, G4 structures may regulate CTCF binding by promoting hypomethylation at consensus DNA motifs. Moreover, the structural basis by which a proximal G4 could influence CTCF binding to its consensus motif remains to be elucidated. Our studies on the other hand provide evidence that CTCF can directly bind G4 structures, corroborating findings from a previous study^24^. We demonstrate that similar to CTCF binding to its consensus DNA motif, G4 recognition by CTCF involves ZFs 4 to 7 (117 amino acids). However, in contrast, a single point mutation (H373R) in the ZF4 of CTCF abolishes binding to the consensus motif while binding to G4s remains unaffected. Moreover, we estimate that the affinity of CTCF for G4s is ∼2-fold lower than its affinity for the consensus motif. Even though our studies imply that the C2H2 motif in ZF4 domain is not required for G4 recognition, the precise structural basis for CTCF-G4 binding is unresolved, and we plan to follow up on this work in future studies.

In another useful contribution, we have developed a novel G4 mapping method that utilizes a highly selective small molecule G4 ligand, NMM^53^. This adds to a growing catalog of G4-mapping methods that each offer unique advantages but may have specific limitations. Notably, G4 mapping using the NMM-Bio method appears to define a set of G4 peaks distinct from those defined by the BG4 antibody. We adapted the NMM-Bio method in the CUT-and-Tag workflow to use with limiting cell numbers. The NMM-Bio G4 mapping strategy can be used to complement existing BG4-based G4 mapping to generate more comprehensive G4 maps for exploring G4 biology.

In addition to the CTCF-G4 interaction, our screen revealed a diverse array of both previously known and novel G4-interacting proteins. Key features of our proteomics screen, distinct from previous approaches, are the use of a mixture of DNA G4s of different topologies and the capture of proteins under native intracellular salt concentrations. These conditions allowed us to detect proteins (such as CTCF) that were not identified by previous G4 proteomics screens. Our work implicates G4 structures in regulation of several genomic processes including RNA splicing, mRNA cleavage, nucleosomal remodeling, histone modifications among others. While many of the hits from our screen were components of multi-subunit protein complexes, the mechanisms by which they interact with G4s remain unknown.

An intriguing category of proteins highlighted in our screen were those associated with membrane-less nuclear bodies with biomolecular condensate-like features such as Cajal bodies and paraspeckles. The enrichment of condensate-associated proteins by G4s raises fascinating questions about the potential roles of G4s in formation of biomolecular condensates, and this is consistent with recent work implicating RNA G4s in liquid-liquid phase separation^60–62^. More broadly, from an evolutionary viewpoint, the number of predicted G4 structures appear to have expanded during hominid evolution when compared across 37 different species^9^. This indicates that G4 structures may have been positively selected during human evolution and co-opted in regulation of diverse genomic processes.

## Acknowledgements

We thank Dr. Elphege Nora for sharing the mESCs carrying the AID-tagged CTCF. We thank Drs. Xinkun Wang and Matthew Schipma and other members of the Northwestern University Sequencing Core for providing next generation sequencing services, the Northwestern University RHLCCC Flow Cytometry Facility team: Dr. Swaminathan S., Ostiguin C., Dias E., and others for their help with cell sorting. Some figure schematics were created using Biorender. This work was supported by the SPARK pilot award, R00 award from the National Cancer Institute (grant ID: CA248835) and institutional startup funds from Northwestern University and the Lurie Cancer center to V.S.

## Author Contributions Statement

DSC, IH, RCM and VS acquired, analyzed and interpreted the data. DSC and VS performed the bioinformatics analysis. SC and BW performed characterization of ESCs. IR helped with designing biochemical assays for CTCF-G4 interaction. AC and FA helped with Hi-C data analysis. SAM performed the MASS-Spec for G4 proteomics screen. AR supervised the initial studies for G4 proteomics screen and biochemical characterization of CTCF-G4 interaction. VS supervised the studies, conceptualized the experiments and helped with data interpretation. DSC, IH, RCM and VS wrote, and all members of the Shukla lab reviewed the manuscript. All authors were involved in reviewing and editing the manuscript.

## Data Availability

All genome-wide sequencing datasets have been deposited to Gene Expression Omnibus (GEO) repository, accession number GSE288430 and GSE288516. Any data and reagents will be made available upon request.

## Code Availability

The code used to process the next-generation sequencing datasets are deposited to github (https://github.com/dsamanie7/G4_CTCF).

## Materials and Methods

### Cell culture

Murine B cells from C57Bl/6, mice were isolated using EasySep Mouse naive B cell isolation kit (#19484 Stem Cell Technology, Canada) from splenocytes. Primary B cells and CH12F3 (CH12) cells were cultured at 37°C, 5% CO_2_ in RPMI 1640 media supplemented with 10% FBS, 1x MEM non-essential amino acids, 10 mM HEPES (pH 7.4), 2 mM Glutamax, 1 mM sodium pyruvate, 55 µM 2-mercaptoethanol (all from Life technologies). B cells (5×10^5^-1×10^6^ cells/mL) were activated with 25 µg/mL LPS from *E. coli* O55:B5 (Sigma, St. Louis, MO) and 10 ng/mL rmIL-4. For pyridostatin (PDS) (#SML2690, Sigma-Aldrich) and RHPS4 (Tocris) treatment, CH12 cells were incubated in drugs for 24 hours before analysis. All the cytokines used above were from Peprotech (Rocky Hill, NJ). The CH12F3 cell line was developed in Dr. T. Honjo’s laboratory at Kyoto University and was independently validated using cell stimulation and Ig class-switching experiments.

E14 and EN52.9.1 (CTCF-AID) ESCs were obtained from Dr. Elphège Nora (University of California, San Francisco). mESCs were cultured at 37°C with 5% CO_2_ in DMEM supplemented with 15% ESC-grade fetal bovine serum (ES cell grade FBS from ThermoFisher), 2 mM GlutaMAX, 1X MEM non-essential amino acids, 1 mM sodium pyruvate, 55 µM β-mercaptoethanol (all from Life Technologies), and 1000 U/mL ESGRO murine leukemia inhibitory factor (mLIF Cat# ESG1106, Millipore Sigma). For auxin-induced degradation of CTCF, media was supplemented with 500 µM auxin analog indole-3-acetic acid sodium salt (IAA, Cat# I5148-2G, Millipore Sigma). Auxin washout was performed by washing cells twice with PBS or DMEM.

### Detection of G-quadruplex binding proteins

B cells activated in presence of 25 µg/ml LPS and 10 ng/ml IL-4 or CH12F3 cells were used to collect nuclear extracts with Dignam extraction. Cells were resuspended in 10 mL Dignam buffer A (10 mM KCl, 1.5 mM MgCl_2_) supplemented with protease inhibitor cocktail (Thermofisher) and incubated on ice for at least 10 min. The cells were then homogenized and lysed with a dounce homogenizer (10 strokes) and nuclei were collected by centrifugation at 600 g for 5 min and washed again with buffer A. Cells were then resuspended in 1ml of Dignam buffer C (0.2 mM EDTA, 25% glycerol (v/v), 20 mM HEPES-KOH (pH 7.9), 0.42 M NaCl, 1.5 mM MgCl_2_) supplemented with protease inhibitor cocktail and rotated at 4°C for 1 hour. The lysates were collected and passed through a PD-10 buffer exchange column (GE Healthcare) equilibrated with Intracellular Buffer (25 mM HEPES pH 7.5, 10.5 mM NaCl, 110 mM KCl, 130 nM CaCl_2_, 1 mM MgCl_2_). The G4 and non-G4 oligonucleotides (**Fig.1A**) were folded in the presence of 10mM HEPES (pH 7.4) and 150mM KCl, annealed with a complementary biotinylated adaptor denatured at 95°C for 5 minutes with temperature ramp down to 25°C at 0.1°C/second. The annealed oligos were then captured on streptavidin conjugated magnetic beads and incubated with nuclear lysates to identify G4-binding proteins. For initial studies, the nuclear lysates collected in Intracellular Buffer were spike-in with 1 µg FLAG-tagged BG4 scFv antibody (Cat# MABE917, Millipore Sigma), pre-cleared with MyOne T1 streptavidin-conjugated DynaBeads (Cat# 65-601, Invitrogen) for 1 hour and incubated with streptavidin beads conjugated with a mixture of G4 or non-G4 forming oligonucleotides for 4 hours. Samples were then washed with Intracellular Buffer with 0.05% Tween-20 for a total of 5 washes and denatured using the LDS-PAGE sample buffer (Cat# NP0008, Invitrogen) for immunoblotting. The oligo sequences are included in **supplementary table 1** and immunoblotting antibodies in **supplementary table 2**.

### MASS-Spectrometry for Proteomics screen

For the proteomics screen, G4 and non-G4 control pull-down fractions were loaded on denaturing gel and stained with silver stain (Silver stain kit from Pierce). Streptavidin beads from the pulldown were resuspended in 50 uL of 50 mM Tris pH 8, 150 mM NaCl, frozen and stored at –80C until processing. For processing, dry urea was added to a final concentration of 8 M and incubated at room temperature for 15 min. Urea was diluted to 4 M with the same buffer and 20 units of LysC (Wako) was added and incubated at 37°C for 60 min. The supernatant was transferred to a new tube at 37°C and the beads were digested further with 250 ng trypsin (Promega) for 60 min. The trypsin digest was combined with the LysC digest, urea was diluted to < 2 M and allowed to digest overnight. Peptide digests were desalted, TMT (Thermo) labeled “on-column”, mixed, and fractionated into six basic reversed phase fractions before being concatenated back into three before analysis by liquid chromatography tandem mass spectrometry (ref: PMC6495249). Liquid chromatography was performed using a Proxeon EASY-nLC 1000 at a flow rate of 200 nl/min. Peptides were separated at 50°C using a 75 micron i.d. PicoFit (New Objective) column packed in house with 1.9 um AQ-C18 material (Dr. Maisch) to 20 cm in length over an 84 min effective gradient. Mass spectrometry was performed on a Thermo Scientific Q Exactive Plus. After a precursor scan from 300 to 2,000 m/z at 70,000 resolution the top 12 most intense multiply charged precursors were selected for HCD at a resolution of 35,000. Data were searched with Spectrum Mill (Agilent) using the Uniprot Mouse database with 261 common laboratory contaminants. A fixed modification of carbamidomethylation of cysteines and variable modifications of N-terminal protein acetylation, oxidation of methionine, and TMT 11-plex labels were searched. The enzyme specificity was set to LysC/ trypsin and a maximum of three missed cleavages was used for searching. The maximum precursor-ion charge state was set to 6. The precursor mass tolerance and MS/MS tolerance were set to 20 ppm. The peptide and protein false discovery rates were set to 1%.

For statistical enrichment analysis, we performed a moderated T-test followed by Benjamini-Hochberg correction for false discovery rate estimation. The enriched samples were the numerator, and the non-G4 DNA sequences were used at the denominator. The positive hits from MASS-Spec are included in **supplementary table 3 and 4.**

### Immunoblotting

Protein lysates were resolved using a NuPAGE 4-12% Bis-Tris gel (Cat# NP0321BOX, Invitrogen) and transferred to PVDF membrane (Cat# IB23001, Invitrogen) using the iBlot 2 Semi-Dry Gel Transfer system (Cat# IB21001, Invitrogen) at 25 V for 7 min. Membrane was blocked with 5% nonfat dry milk in TBS-T buffer (1 M Tris-HCl pH 7.4, 5 M NaCl, 2.5 M KCl, 0.5 M EDTA, 0.05% Tween-20) and incubated with an appropriate primary antibody (**supplementary table 2**), followed by incubation with horseradish peroxidase (HRP)-conjugated secondary antibody, diluted in 5% milk solution. Each antibody incubation was followed by three 10-min washes with TBSTE (50mM Tris-HCl pH 7.4, 150mM NaCl, 0.05% Tween-20, 1mM EDTA) buffer at room temperature. Signal was detected with enhanced chemiluminescence reagents (Invitrogen) with either X-ray film or imaging with Chemidoc Luminescence system (BioRad). The antibodies for immunoblotting are included in **supplementary table 2**.

### Site-directed mutagenesis of CTCF

Site-directed mutagenesis to introduce H373R mutation in CTCF was performed following the Q5 Site-Directed Mutagenesis Kit from New England BioLabs (NEB Cat# E0554S) with modifications. Forward and reverse primers designed for the mutation were synthesized by Integrated DNA Technologies (IDT). Template plasmid (10 µg) was mixed with forward and reverse primers (each 10 µM) and Q5 Hot Start High-Fidelity 2X Master Mix (NEB Cat# M0494S) for amplification. Following the PCR amplification, the product was subjected to a Kinase, Ligase, and Dpnl (KLD) reaction at room temperature for 5 minutes. Subsequently, the KLD reaction product was transformed into chemically competent *E*. *coli* cells (NEB Cat# C2987H). Individual colonies were picked and cultured overnight in LB medium with kanamycin at 37 °C, and plasmid DNA was extracted using PureLink Quick Plasmid Miniprep Kit (Invitrogen Cat# K210010). The final plasmid DNA was sequenced to confirm the successful introduction of mutation. Sequencing was performed by ACGT DNA Sequencing Services.

### *In vitro* translated CTCF

*In vitro* transcription and translation of CTCF were performed using the TNT Coupled Reticulocyte Lysate System (Promega Cat# L4610) according to the manufacturer’s instructions. Briefly, 1 µg of plasmid DNA template with CTCF open reading frame downstream to the T7 promoter was combined with TNT reaction buffer (Promega Cat# L462A), TNT T7 polymerase (Promega Cat# L481A), amino acid mixture (Promega Cat# L995A-L996A), TNT rabbit reticulocyte lysate (Promega Cat# L483A), and recombinant RNase inhibitor (Takara Bio Cat# 2313A). The reaction mixture was incubated at 30°C for 90 minutes on a thermocycler. A small portion of the final protein product (∼2 µL) was kept for immunoblotting as input, and the remaining product was directly incubated with annealed biotinylated G4 oligos to assess G4-CTCF binding (please see below, CTCF-G4 binding assay).

### Recombinant CTCF purification

Coding sequences of N-terminal Flag-tagged human CTCF fragment ZF4-7 WT and H373R mutant (amino acid 350-466) were sub-cloned in frame with Glutathione-S-Transferase (GST) protein separated by a PreScission protease cleavage site in the pGEX6p1 vector. BL21 (DE3) cells were transformed with the pGEX6p1 vector with GST-CTCF ZF4-7 fusion. Single colonies were picked and grown overnight in 5 mL LB at 37°C before further expanding in 2 to 3 litres of LB (1:100) with shaking at 37°C for 2-3 hours until the OD at 600 nm reached 0.5-0.6 (log phase). ZnCl_2_ was added to a final concentration of 25 μM, expression was induced by the addition of isopropyl β-D-1-thiogalactopyranoside (IPTG) to 0.5 mM, and the cultures were incubated overnight at 16°C. Cells were harvested by centrifugation, resuspended in lysis buffer containing 20 mM Tris-HCl (pH 7.5), 500 mM NaCl, 5% (v/v) glycerol, 0.5 mM Tris (2-carboxyethyl)-phosphine hydrochloride (TCEP), and 25 μM ZnCl_2_. The cell suspension was then lysed by sonication for a total of 4 min (5 seconds on, 30 seconds off cycle) with a probe sonicator (Missonix) followed by centrifuge at 10,000 g for 20 min, at 4°C. Lysates were mixed with polyethylenimine (Sigma) at pH 7 to a final concentration of 0.3%–0.4% (w/v) before centrifugation at 16,500 rpm. Expression of protein was confirmed by Coomassie staining on SDS-PAGE gel by comparing with an uninduced control culture. The supernatant was then mixed with 2ml washed and equilibrated glutathione magnetic beads (Cat# 78602, Pierce) and rotated for 4 hours overnight. The lysate beads mixture was then washed three times with lysis buffer and then re-suspended in 2 mL elution buffer (100 mM Tris-HCl (pH 7.6), 10% Glycerol, 25 μM ZnCl_2_, 100 mM KCl, 1 mM DTT) and treated overnight with PreScission protease (Cat# Z02799, Genescript). The supernatant was then collected, and the proteins were purified by gel filtration on a Superdex 200 column (GE Healthcare). The protein concentrations were quantified by absorbance at 280 nm using Nanodrop. The purity of protein was additionally confirmed by Coomassie staining of samples on an SDS-PAGE gel.

### CTCF binding to G4 and consensus motif

To measure CTCF-G4 binding, *in vitro*-translated or recombinant CTCF protein were incubated with either a mixture of three different biotinylated G4 (*Kit2, Kit1 and Spb1*) and non-G4 oligos, all of which showed similar binding to CTCF (**Fig. 1G**). For quantitative binding assays, the *Kit2* G4 or non-G4 control oligos were used. For CTCF binding to consensus DNA motif, the double stranded CTCF consensus or mutated consensus oligos were annealed to the same biotinylated adaptor also used for G4 binding assays. The CTCF incubation with different oligos was performed in Intracellular Buffer (25 mM HEPES pH 7.5, 10.5 mM NaCl, 110 mM KCl, 130 nM CaCl_2_, 1 mM MgCl_2_) containing 0.125% BSA for 1 hour at 4°C with rotation. MyOne T1 streptavidin-conjugated DynaBeads (Cat# 65-601, Invitrogen) beads were pre-washed five times with Intracellular Buffer containing 0.125% BSA and were used to capture biotinylated oligos at 4°C for 1 hour with rotation. After incubation, samples were washed four times with Intracellular Buffer containing 0.05% Tween-20. To reveal the signal in some experiments, the binding was assessed by immunoblotting. In other cases, for quantitative binding measurements, signal was measured by on-bead fluorescence with flow cytometry (Cytek). For immunoblotting based measurements, the western blotting under denaturing conditions were performed as described above (**Antibodies, supplementary table 2**). For competition assays in **Fig. 2D**, the non-biotinylated double stranded CTCF consensus or mutated consensus oligos were mixed at different molar concentration with G4 or non-G4 oligos.

For quantitative binding assays, streptavidin-conjugated magnetic beads (Invitrogen, 65-601) with captured oligos were incubated with anti-Flag primary antibody (Cat# 14793S, 1:100 dilution, Cell Signaling) for 30 min at 4°C with rotation, followed by two washes and a 30-min incubation with a PE-conjugated anti-rabbit secondary antibody (Cat# 406421, 1:400 dilution, Biolegend) at 4°C with rotation. Beads were washed, resuspended in intracellular Buffer, and on-bead fluorescence was measured using flow cytometry (Cytek). For the detection of binding between CTCF and consensus motifs, the G4 or non-G4 oligos were replaced with CTCF oligos (**supplementary table 2**). For comparison with the conditions used in the study by Wulfridge et al.^20^, Intracellular Buffer in this assay was replaced with Binding Buffer (50 mM TRIS pH 8.0, 100 mM NaCl, 1 mM DTT, 0.1 mM EDTA, 5% glycerol, 0.05% NP-40). Additional assays were performed under identical conditions with EDTA removed to assess its impact on CTCF-G4 interaction.

### NMM-Bio conjugation and purification

The NMM-Biotin conjugation was performed by a commercial vendor WuXi AppTec. 5mg NMM (8.61 μmol) was dissolved in anhydrous Dichloromethane, DCM (2 mL). Under nitrogen the NMM solution was added to 2 molar equivalents of Propylphosphonic anhydride (T3P, 13.21 mg, 20.75 μmol), 5 molar equivalents of N,N-Diisopropylethylamine (DIEA, 6.71 mg, 51.88 μmol) and Amine-PEG11-Biotin (8 mg, 10.38 μmol) at 20°C. The reaction was stirred at 20°C for 12 hours. After that, thin layer chromatography (TLC) with DCM/Methanol = 9/1, stained by visible light was performed and showed complete consumption of NMM (retention factor = 0.6) in the reaction. TLC (DCM/Methanol = 4/1, stained by visible light) also showed complete consumption of Amine-PEG11-Biotin (Retention factor= 0.3) in the reaction. Seven similar scale reactions were performed as described above and combined to generate a large batch of NMM-Bio. The reactions were concentrated under reduced pressure at 20 °C to remove extra DCM. After full wavelength scanning, the NMM-Bio complex showed maximum absorption at 395 nm, as also described earlier^53^. Following this, NMM-Bio was purified by a silica column under 395 nm wavelength with medium pressure liquid chromatography (MPLC) using a Welch Ultimate XB NH2 column with Heptane and Ethanol. The NMM-Bio conjugation was confirmed by 1H-NMR and high-resolution mass spectrometry (HRMS-TOF). Finally, the NMM-Bio complex was concentrated and lyophilized (21.5 mg, 10.3 μmol, 14.2% yield) as deep red color solid.

### CUT-and-Tag library preparation, sequencing, and analysis

The CUT-and-Tag protocol from EpiCypher was adapted with modifications. Approximately 300,000 ESCs were collected from 24 hour auxin treatment. The isolated nuclei from mESCs were mixed with activated Concanavalin A (ConA)-coated magnetic beads (Cat# 21-1401, Epicypher) and incubated for 10 minutes at room temperature. Upon nuclei extraction and binding nuclei to activated beads, incubation steps were completed in a modified Digitonin 150 buffer (20 mM HEPES pH 7.5, 150 mM KCl, 0.5 mM spermidine, 1x protease/phosphatase inhibitor) until the tagmentation step. This modification, in which NaCl was replaced by similar concentration of KCl, was to optimize stabilization of G4 structures during prolonged incubation steps, as shown previously^13^. Other buffers were prepared and used according to manufacturer’s protocol. The nuclei-conjugated ConA bead complexes were incubated overnight at 4 °C with primary antibody (CTCF, BG4, NMM-Bio, or control NMM; **supplementary table 2**) in the modified Digitonin 150 buffer. After primary incubation, nuclei-bead complexes were incubated with secondary anti-biotin antibody (Cat# ab53494, Abcam) in the modified Digitonin 150 buffer for 1 hour at 25 °C, followed by tagmentation using CUTANA pAG-Tn5 (Cat# 15-1117, Epicypher). Targeted chromatin tagmented products were column purified using a DNA Clean & Concentrator Kit (Cat# D4013, Zymo Research). DNA libraries were amplified with the following thermocycling conditions: initial denaturation at 98°C for 30 seconds, 18 cycles of denaturation at 98 °C for 10 seconds, annealing at 63°C for 30 seconds, and extension at 72 °C for 1 minute, followed by a final extension at 72 °C for 1 minutes. After PCR amplification, the libraries were column purified once again to be quantified and qualified before pooling for sequencing. Purified libraries were sequenced on Illumina NovaSeq 6000.

Paired-end reads (50bp) were mapped to the mouse genome mm10 GRCm38 (Dec. 2011) from UCSC and E.coli K-12 genome (NZ_CP132594.1) with Bowtie (“--p 8)^63^. Reads mapping to the *E.coli* genome were filtered out from sam files using ‘grep’ command in bash. The remaining mm10 mapped reads were marked for duplication and kept with picard-tools-2.21.4 MarkDuplicates (https://broadinstitute.github.io/picard/). Peaks were called using MACS2 (2.1.4) (“callpeak –-keep-dup all” using IgG samples as input)^64^. The intersected peaks from the same condition were obtained using HOMER^65^ mergePeaks and filtered with grep. Intersection between the different methods were performed with bedtools^66^ “intersect” (bedtools/2.29.2) or HOMER mergePeaks. Deeptools^67^ (3.5.5) was used to plot the signal at the TSS and center of the peaks.

### Lentiviral production and transduction of mESCs

HEK293T cells were cultured to 80% confluence and transfected using the Lipofectamine® 3000 Transfection Kit (Invitrogen, Cat#L3000001), following manufacturer’s protocol, with 2.5µg DNA (1.25µg lentiviral plasmid; 3-parts psPAX2, 1-part pMD2.G – and 1.25µg EV, FL or H373R plasmids). Media containing lentivirus secreted by HEK293T cells was collected after 48 hours and transfection was assessed by flow cytometry for mCherry. Media was aspirated from ESCs cultured in a 6-well plate and replaced with a 1:1 mix of lentiviral media and DMEM ESC media (0.5 multiplicity of infection) with polybrene (10ng/ml) and centrifuged at 2000 rpm for 90 minutes at 37°C to facilitate transduction. Following transduction, cells were rested for 4 hours before aspirating viral media and replacing with fresh DMEM. Transduced cells were sorted by fluorescence-associated cell sorting based on expression of mCherry reporter. Sorted cells were expanded in culture for one week in DMEM ESC media and assessed again for mCherry expression prior to conducting experiments.

### RNA-seq library preparation, sequencing, and analysis

RNA was purified from cell lines treated with or without auxin using the RNeasy Plus Mini Kit (Qiagen) following manufacturer’s instructions. Purified RNA (200 ng) was subsequently utilized to prepare RNA libraries with the NEBNext Ultra II Directional RNA Library Prep Kit (New England Biolabs, Cat# E7760), following manufacturer’s recommended protocol. Samples were sequenced on the Illumina Novaseq 6000 platform. Paired-end (50 bp) reads were mapped to the mouse genome mm10/GRCm38 using STAR4 (2.7.10a)^68^ (--genomeLoad LoadAndRemove –-outFilterMismatchNmax 4 –-outFilterMultimapNmax 100 –-winAnchorMultimapNmax 100). Counts were obtained with featureCounts^69^ (subread v2.0.3)5 –p –B –g gene_name –s 2 –-countReadPairs). Differentially expressed genes were calculated with DESeq2^70^, filtering out genes that did not have any count in any condition.DEGs were defined using an adjusted p-value≤0.05 and a log2 fold change ≥±0.58. Pathway enrichment analysis of DEGs was performed using Metascape (https://metascape.org). TC-seq package^71^ was used on RNA-seq read counts to detect the temporal pattern of the differential gene expressions (RPKM values) (“algo = ‘cm’, k = 9). DEGs were assigned to a cluster (C1-C9) representing a specific temporal pattern of expression.

### Motif analysis

For de novo and known motif scanning, we used HOMER “findMotifsGenome.pl” function to identify de novo and known motifs in the different subsets of CTCF peaks. In parallel, we also performed motif discovery analysis with MEME (5.5.2) with the following parameters: “-dna –nmotifs 20 –minw 6 –maxw 50 –revcomp”. For identifying putative G4 forming motifs, we used regular expressions in ‘awk’commands to categorize potential G4 forming mitifs from fasta sequences. The loop1–7 was defined with the following expression: G_3_ N_1–7_ G_3_ N_1–7_ G_3_ N_1–7_G_3_. The G4s with long loops were defined as sequences with any loop length >7 (up to 12 for any loop and 21 for the middle loop). The simple bulge was defined as sequences with a G4s with a bulge of 1–7 bases in one G-run or multiple 1-base bulges. The two tetrad/ complex bulge was defined as sequences with G4s with two G-bases per G-run with several bulges of 1–5 bases.

### Hi-C analysis

We downloaded all available Hi-C replicates for Untreated, auxin-inducible degron (AID), and Washoff conditions from the GEO repository (GSE98671) to investigate chromatin interactions under different treatments. The raw Hi-C reads were processed using the HiC-Pro v2.8.0 pipeline^72^, which involved mapping the data to the mm10 genome assembly to generate genomewide interaction matrices. To identify significant chromatin interactions, we employed the Fit-Hi-C v1.1.3 tool^73^, configuring it with the options “-U 5000000” and “-L 20000” to call interactions at a 20Kb resolution. This approach allowed us to robustly detect and analyze significant genomic interactions across the various experimental conditions. We further filtered the Hi-C interactions by selecting only those with a q-value < 0.05.

---

**Supplementary Figure 1.**
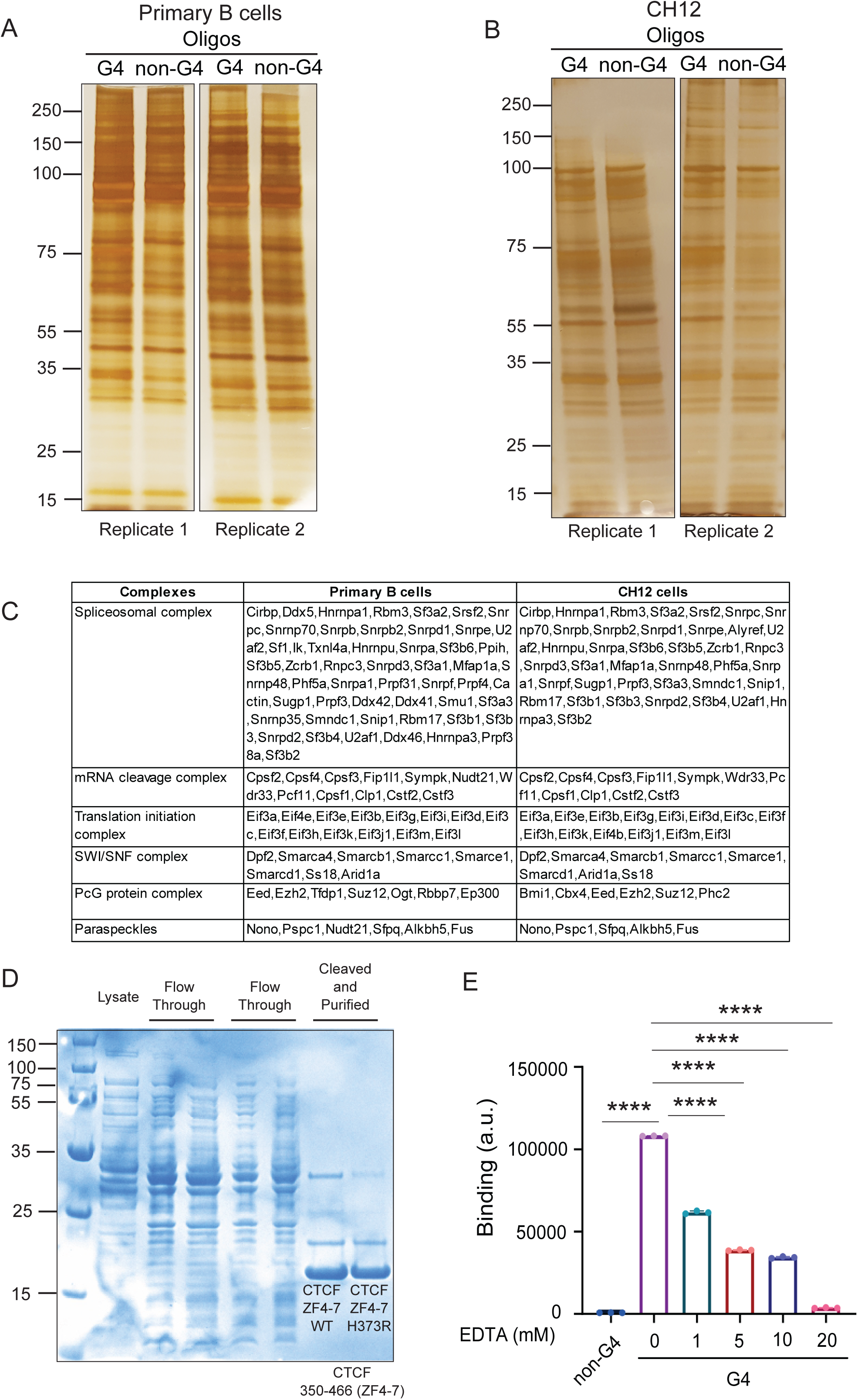
A. Silver stain of pulldown fractions from G4 and non-G4 control oligos in the nuclear lysates of LPS and IL-4-stimulated primary B cells. B. Silver stain of pull-down fractions from G4 and non-G4 control oligos in the nuclear lysates of CH12 cells. C. List of protein complex subunits enriched in G4 proteomics screen of primary B cells and CH12 B cells. Multiple subunits within each complex were enriched. D. Expression and purification of recombinant CTCF ZF4-7 WT or H373R (ZF4-7) mutant from *E. coli*. E. Binding assay showing the interaction of recombinant CTCF ZF4-7 protein with G4 (red, circle) or non-G4 (blue, square) oligos in the absence or presence of increasing concentrations of EDTA. X-axis denotes CTCF protein concentrations in nanomolar (nM), Y-axis represents arbitrary units (a.u.) of binding. **** p value ≤ 0.0001 calculated with student T-test.

**Supplementary Figure 2.**
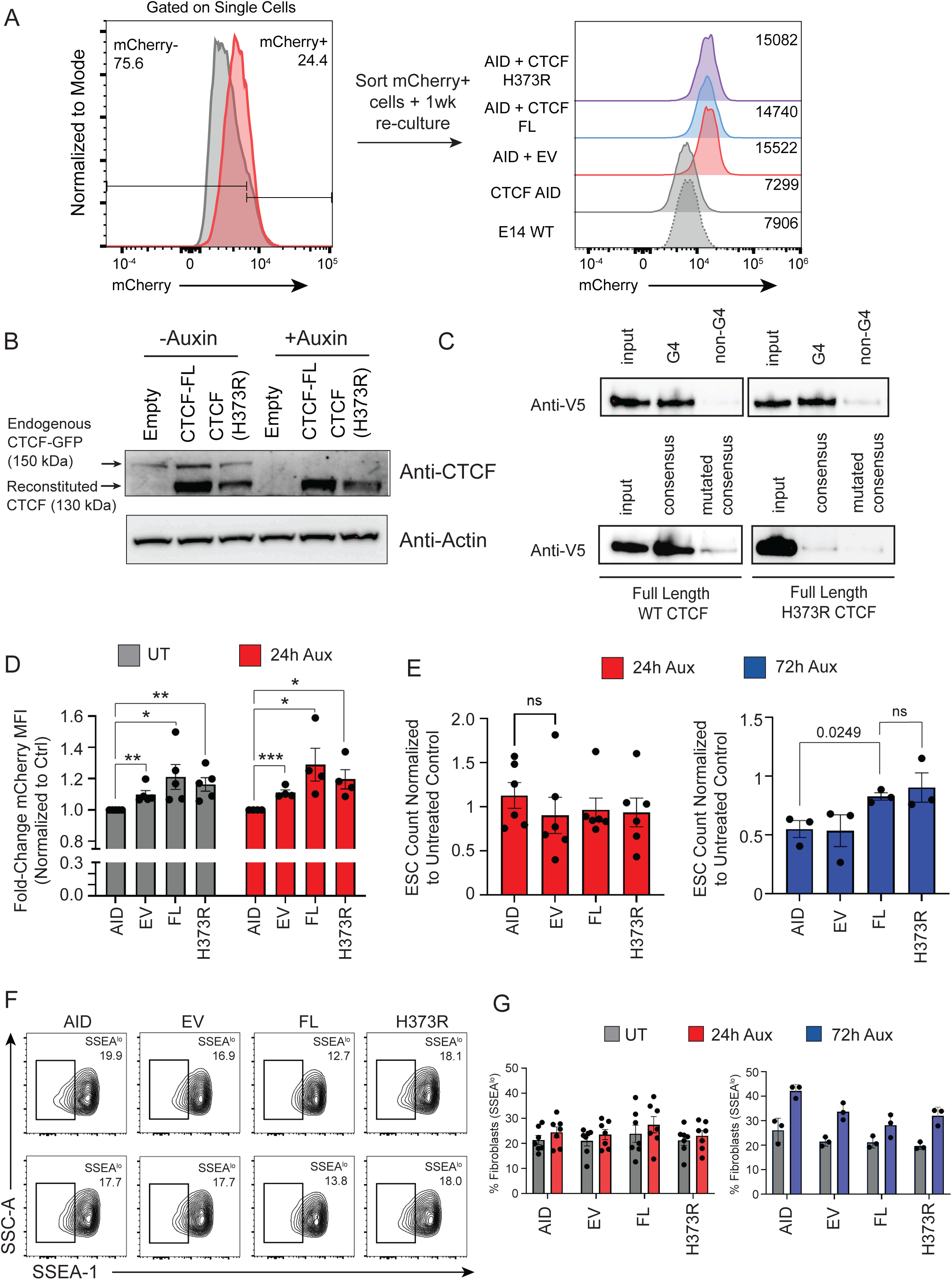
A. Sorting strategy for mESCs harboring AID-tagged CTCF transduced with empty vector (EV), CTCF FL and CTCF H373R constructs expressing mCherry reporter. mCherry+ cells were sorted and expanded in culture for one week prior to CUT-and-Tag. B. Western blot for CTCF expression in cell lysates derived from EV, FL and H373R expressing mESCs after 8 hours with or without auxin treatment. Beta-Actin was used as a loading control. C. Immunoblot showing the enrichment of *in vitro* transcribed and translated full length or H373R CTCF pulled down with either G4, non-G4, CTCF consensus, or mutated consensus oligos. D. Barplots quantifying the median fluorescence intensity (MFI) of mCherry in AID, EV, FL and H373R expressing mESCs after 24h with and without auxin treatment. mCherry MFI was normalized to the non-transduced AID-CTCF (AID) mESCs in the same experiment. * p value ≤ 0.05, ** p value ≤ 0.01, *** p value ≤ 0.001, calculated using Student T-test. E. Barplots quantifying normalized cell counts in mESCs expressing EV, FL and H373R with auxin treatment for 24 and 72 hours. Cell counts were normalized to the respective untreated controls in each condition. The ratio below 1 indicates reduced cell numbers. Marked p values calculated using Student T-test. F. Representative flow cytometry plots for SSEA-1 expression in AID, EV, FL and H373R mESCs 24 hours with or without auxin treatment. G. Barplots quantifying the frequencies of fibroblasts (SSEA^lo^) in AID, EV, FL and H373R mESCs at 24 and 72 hours with or without (untreated, UT) auxin treatment.

**Supplementary Figure 3.**
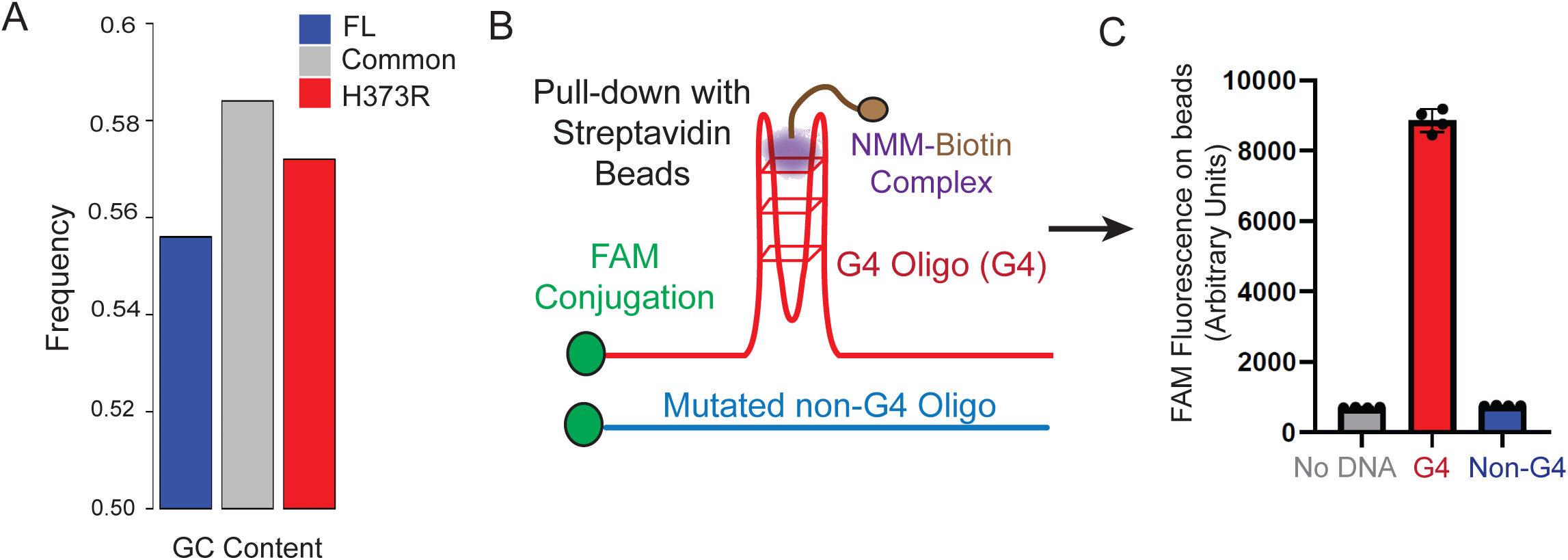
A. Barplot showing GC content across FL only (blue), common (grey) and H373R only (red) CTCF peaks. B. Schematic representation of the experimental design of NMM-Bio binding to a G4 oligo. C. A fluorescein (FAM)-conjugated *Kit2* G4 oligo (red) or mutated non-G4 control (blue) was incubated with NMM-Bio and captured on streptavidin beads. The fluorescence signal was measured using flow cytometry. Data is representative of 3 independent replicates.

**Supplementary Figure 4.**
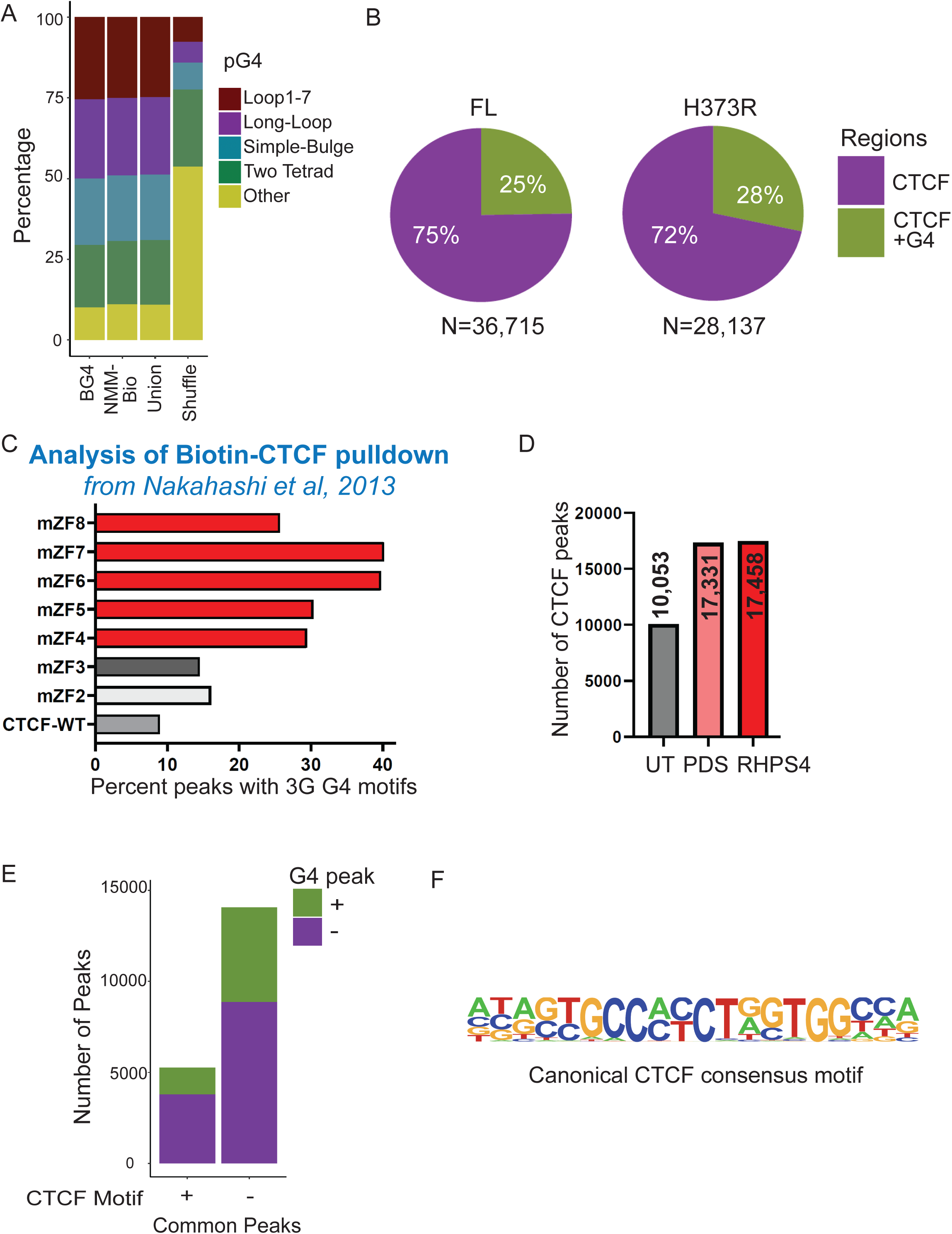
A. Barplot showing different classes of predicted G4 motifs in BG4, NMM-Bio, or union of BG4 and NMM-Bio peaks that overlap regions identified in G4-seq. Genomic shuffle regions are used as controls. Please also see methods for details about the respective G4 motifs. B. Pie-charts showing the percent of CTCF peaks that overlap with G4 peaks (green) identified in mESCs expressing FL or H373R CTCF. C. Bargraph quantifying percent CTCF peaks with 3G-G4 motifs (G_3_N_1-7_G_3_N_1-7_G_3_N_1-7_G_3_). CTCF ChIP-seq was generated with BirA-tagged wildtype (WT) and ZF mutant CTCF proteins overexpressed in B cells^39^. Each ZF from ZF2 to ZF8 was mutated at the second histidine in the C2H2 motif. D. Quantification of the number of CTCF peaks identified using ChIP–seq in B cell line CH12 cells treated with or without G-quadruplex stabilizing ligands PDS (10μM) and RHPS4 (2μM) for 24 hours. E. Bar plot showing the number of common (FL and H373R) CTCF peaks separated based on the presence (+) or absence (-) of a canonical CTCF motif. The peaks were further categorized based on their overlap with (+) or without (-) a G4 peak. F. A weighted matrix of the canonical CTCF motif used in the analysis in panel E. and Figure 5.

**Supplementary Figure 5.**
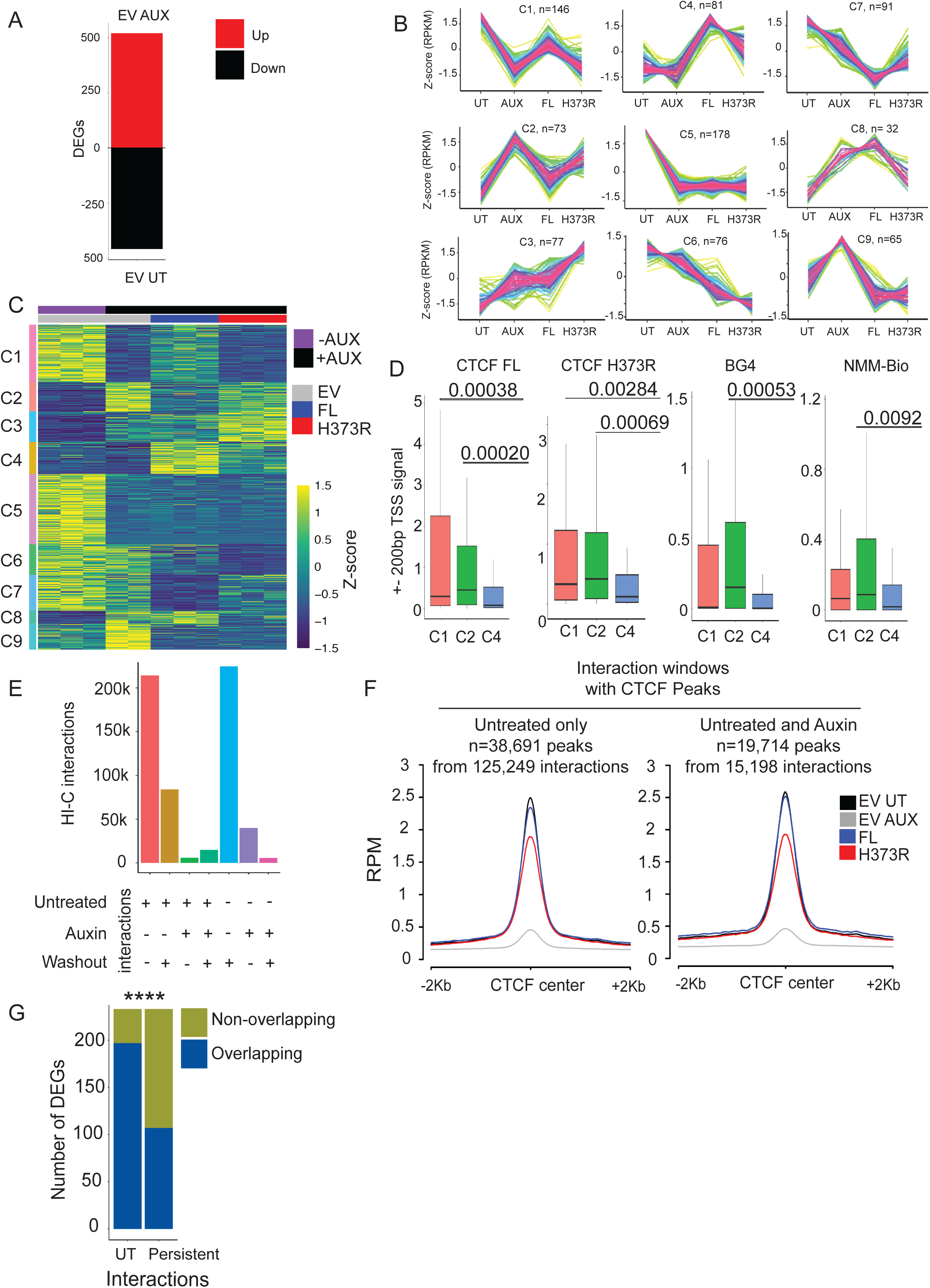
A. Quantification of the differentially expressed genes in EV-AUX vs EV-UT pair-wise comparison. Genes are separated into up (red) and downregulated (black) B. Analysis of temporal gene expression modules with TC-seq. Gene expression profiles were analyzed in EV-UT, EV-AUX, FL and H373R cells and genes were clustered based on their temporal expression patterns. Nine clusters (C1-C9), or expression patterns, were identified. Y-axis indicates the Z-score calculated using RPKM. X-axis indicates the sample condition. C. Heatmap showing the Z-score from the RPKM expression of EV-UT, EV-AUX, FL and H373R in the nine clusters (C1-C9) identified in TC-seq. D. Box and whisker plots comparing the signal of CTCF FL, CTCF H373R, BG4 FL and NMM-Bio within ± 200bp of the TSS of genes in clusters C1, C2, and C4. The statistical significance was calculated using Wilcoxon signed-rank test. E. Barplot reporting the total number of Hi-C interactions in untreated (UT), auxin and washout conditions. The conditions are represented as + or – to highlight conditions in which the Hi-C interactions were identified. F. Histograms displaying CTCF signal from EV-UT, EV-AUX, FL and H373R ESCs plotted in Hi-C interactions with windows overlapping CTCF peaks in untreated or persistent interactions in untreated and auxin treated conditions. G. Barplot showing the total number of DEGs (promoters) that overlap (blue) or do not overlap (non-overlapping; olive) Hi-C interactions from untreated (UT) condition or those that persist (persistent) upon auxin treated conditions. The statistical significance is calculated using Chi-square test, **** p-value <0.00001.

**Supplementary Figure 6.**
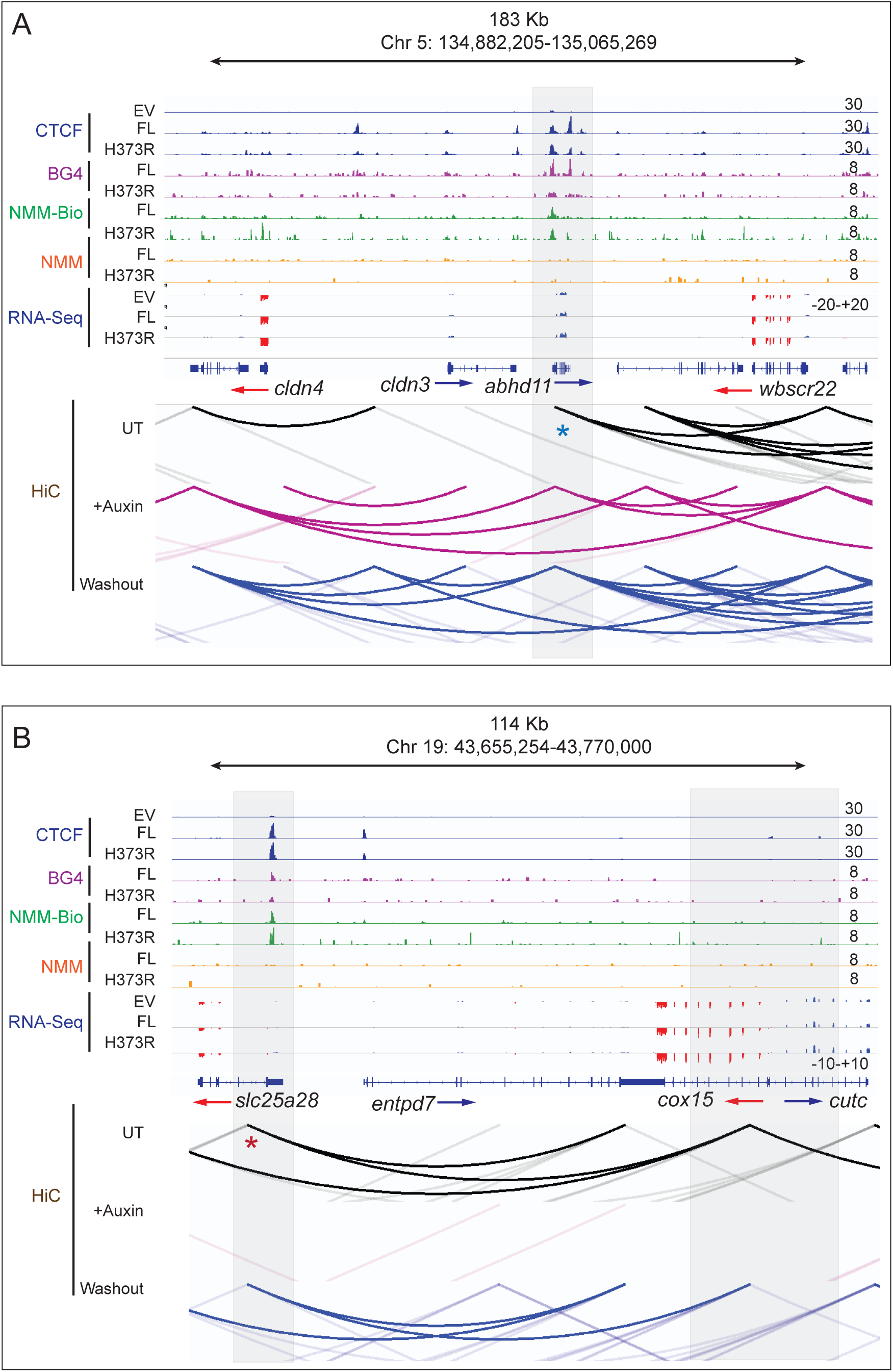
**A and B**. Representative genome browser view of CTCF (blue), BG4 (purple), NMM-Bio (Green), NMM (Orange) and RNA-seq (blue/red) at *Cldn3* (A) and *Cox15* (B) gene loci. Shaded areas define regions of interest with CTCF and G4 peaks. Bottom part is HiC contact maps in the parent mESCs (AID-tagged CTF) without auxin (untreated, UT), with auxin treatment for 2 days (+Auxin) and 2 days following auxin washout (Washout). The red and blue asterisks highlight the interaction anchor points that are lost or retained upon auxin treatment.

