## Supplementary_table_1 for "CTCF directly binds G-quadruplex structures to regulate genome topology and gene expression"

| **Oligonucleotides** | **All 5’ to 3’** |
| --- | --- |
| *Complementary ssDNA* | ATGTATCTACTCGCAGTATCATAC-**3’BiosG** |
| *Kit1* | GATACTGCGAGTAGATACAT*GAAGGGAGGGCGCTGGGAGGAGGGATCT* |
| *Kit2* | GATACTGCGAGTAGATACAT*GAACGGGCGGGCGCGAGGGAGGGGATCT* |
| *Spb1* | GATACTGCGAGTAGATACAT*GAAGGCGAGGAGGGGCGTGGCCGGCATCT* |
| *Telo* | GATACTGCGAGTAGATACAT*GAATTAGGGTTAGGGTTAGGGTTAGGGATCT* |
| *Kit1 non-G4* | GATACTGCGAGTAGATACAT*GAAG****CA****AG****CA****CGCTG****CA****AGGAG****CA****ATCT* |
| *Kit2 non-G4* | GATACTGCGAGTAGATACAT*GAACG****CA****CG****CA****CGCGAG****CA****AG****CA****GATCT* |
| *Spb1 non-G4* | GATACTGCGAGTAGATACAT*GAAGGCGA****CA****AGG****CA****CGTGGCC****CA****CATCT* |
| *Telo non-G4* | GATACTGCGAGTAGATACAT*GAATTA****CA****GTTA****CA****GTTA****CA****GTTA****CA****GATCT* |
| *CTCF Consensus (****Fig.2C****)* | GGCCAGCAGGGGGCGC |
| *CTCF Consensus Mutated (****Fig.2C****)* | GTCAATCAGAGTACGC |
| *CTCF Consensus Bio (****Fig.2H-J, S2B****)* | GATACTGCGAGTAGATACAT*GAA*GGCCAGCAGGGGGCGC |
| *CTCF Consensus Mutated Bio*  *(****Fig.2H-J, S2B****)* | GATACTGCGAGTAGATACAT*GAA*GTCAATCAGAGTACGC |
| *Kit2 Fam* | ***Fam5’****-CGGGCGGGCGCGAGGGAGGGGAT* |
| *Kit2 control Fam* | ***Fam5’****-CG****CA****CG****CA****CGCGAG****CA****AG****CA****GAT* |
