## Supplementary_table_2 for "CTCF directly binds G-quadruplex structures to regulate genome topology and gene expression"

| **Method** | **Target/agent** | **Concentration** | **Manufacturer** | **Catalog Number** |
| --- | --- | --- | --- | --- |
| Immunoblotting | FLAG tag | 1:500 | Cell Signaling Technology | 14793S |
|  | BLM | 1:1000 | Santa Cruz Biotechnology | sc-365753 |
|  | THYN1 | 1:1000 | Atlas Antibodies | HPA038732 |
|  | ARID1A | 1:500 | Cell Signaling Technology | 12354S |
|  | SMARCB1 | 1:1000 | Cell Signaling Technology | 91735S |
|  | TRIM28 | 1:1000 | Cell signaling Technology | 4123 |
|  | NONO | 1:1000 | Cell Signaling Technology | 90336S |
|  | CTCF | 1:100 | EMD Millipore | 07-729 |
|  | V5 tag | 1:500 or 1:100 | Cell Signaling Technology | 13202 |
|  | ß-actin | 1:5000 | Cell Signaling Technology | 13E5 |
|  | HRP-linked Rabbit IgG | 1:2000 | Cell Signaling Technology | 7074 |
| CUT&Tag | CTCF | 1:50 | EMD Millipore | 07-729 |
|  | scFv-BG4 | 1:20 | Millipore Sigma | MABE917 |
|  | NMM | 20 uM | Frontier Specialty Chemicals | NMM580 |
|  | In-house biotinylated NMM | 20 uM | N/A | N/A |
|  | Biotin | 1:100 | Abcam | ab53494 |
